# *Larvaworld* : A behavioral simulation and analysis platform for *Drosophila* larva

**DOI:** 10.1101/2025.06.15.659765

**Authors:** Panagiotis Sakagiannis, Hannes Rapp, Tihana Jovanic, Martin Paul Nawrot

## Abstract

Behavioral modeling supports theory building and evaluation across disciplines. Leveraging advances in motion-tracking and computational tools, we present a virtual laboratory for Drosophila larvae that integrates agent-based modeling with multiscale neural control and supports analysis of both simulated and experimental data. Virtual larvae are implemented as 2D agents capable of realistic locomotion, guided by multimodal sensory input and constrained by a dynamic energy-budget model that balances exploration and exploitation. Each agent is organized as a hierarchical, behavior-based control system comprising three layers: low-level locomotion, optionally incorporating neuromechanical models; mid-level sensory processing; and high-level behavioral adaptation. Neural control models can range from simple linear transfer models to rate-based or spiking neural network models, e.g. to accomodate associative learning. Simulations operate across sub-millisecond neuronal dynamics, sub-second closed-loop behavior, and circadian-scale metabolic regulation. Users can configure both larval models and virtual environments, including sensory landscapes, nutrient sources, and physical arenas. Real-time visualization is integrated into the simulation and analysis pipeline, which also allows for standardized processing of motion-tracking data from real experiments. Distributed as an open-source Python package, the platform includes tutorial experiments to support accessibility, customization, and use in both research and education.

**Author summary:** *Larvaworld* was developed to address two key challenges in behavioral neuroscience and computational modeling. First, it responds to the growing call for closer collaboration between experimentalists and modelers by providing a shared platform -a virtual laboratory-where experimental data analysis and behavioral modeling can be seamlessly integrated. By standardizing dataset formats and ensuring identical, unbiased analysis pipelines for experimental and simulated data, *Larvaworld* facilitates methodological consistency and enables rigorous model evaluation.

Second, it aims to bridge a long-standing gap in theory building and computational modeling at the level of the individual behaving organism. Historically, neuroscience has focused on sub-individual processes, while ecology has concentrated on supra-individual dynamics, resulting in discontinuities among the respective modeling approaches. Recent advances, however, have begun to align these fields, with neuroscience incorporating slower homeostatic processes and ecology integrating faster neurally-mediated mechanisms. *Larvaworld* boosts this convergence by adopting a nested, multi-timescale modeling approach, thus achieving behavioral regulation within the normative homeostatic constraints as these dynamically unfold during larval development. By combining established modeling paradigms from neuroscience and ecology, it provides a novel and flexible platform for studying behavior at the level of the individual organism, promoting cross-disciplinary insights and advancing computational neuroethology [1].

## Introduction

*Drosophila* is a widely studied model organism in neuroscience, alongside lamprey, zebrafish, mouse, rat, and monkey, listed in order of increasing nervous system complexity. Researchers working with mammals are acutely aware of the scarcity and high cost of experimental animals, making computational modeling a valuable alternative to reallife experiments. Insects, on the contrary, are more abundant, affordable, and easier to rear, and their use is subject to fewer ethical restrictions. Nevertheless, growing concerns about animal welfare have recently fueled support for alternative research and educational tools aimed at reducing reliance on live animal experiments. Behavioral studies, in particular, often require far larger sample sizes than neurophysiological experiments, further highlighting the need for computational approaches. In this context, virtual laboratories and simulation platforms are emerging as indispensable tools for both scientific research and education [1, 2]. Many academic institutions have already developed and integrated such tools into their curricula, with notable applications in fields like Mendelian genetics and beyond.

Not all scientific fields readily translate to the virtual domain, and behavioral science is particularly challenging in this regard. The widespread use of arbitrary behavioral metrics across different laboratories further complicates experiment replication. Despite significant advances in experimental setups, genetic manipulations, and behavioral recordings, tools for behavioral modeling remain limited, often lacking key functionalities. In particular, tools that support direct comparisons between simulated and experimental datasets are still underdeveloped.

A recent perspective paper, authored by pioneering researchers in AI introduced the concept of the *embodied Turing test* as a potential goal for computational modeling of animal behavior [3]. This approach seeks to bridge neuroscience and AI by generating robotic or virtual animats, whose embodied behavior is indistinguishable from that of real animals. Achieving this goal requires a platform that facilitates continuous and unbiased comparisons between experimental and simulated datasets to more effectively construct, calibrate, and evaluate models.

Here we propose one such tool for *Drosophila* larvae. The core concept is to create and simulate virtual larvae whose behavior can be analyzed using the exact same pipelines applied to real animals. From the platform’s perspective, experimental and simulated datasets are indistinguishable. Experimental data can be imported and automatically converted into a standardized format identical to that produced by the simulations.

To enhance replicability and accommodate variability in behavioral metrics, the platform relies solely on the originally tracked x-y coordinate time series. All additional behavioral metrics are transparently defined and consistently derived from these coordinates for both experimental and simulated datasets. Virtual larvae are modeled with a simplified 2D body structure that mirrors key features of the real animal. Their movements are tracked during simulations as if observed by a virtual tracker, navigating a detailed virtual replica of the real experimental arena.

Once a trackable body is established, behavioral control can be implemented through computational modeling. To maximize flexibility and support integration and expansion, *Larvaworld* adopts a hybrid, modular, and hierarchical control architecture [4]. Its hierarchical design follows the layered behavioral control paradigm [5], in which successive control layers are stacked according to two key principles: decentralization and subsumption. Decentralization ensures that each layer can operate autonomously, independent of top-down input-consistent with evidence of decentralized neural circuits capable of generating simple behavioral primitives even when isolated from higher-order control [6]. Subsumption, on the other hand, allows top-down modulation to influence only a few key parameters, reflecting subtle adjustments by higher neural centers.

Each control layer is composed of interconnected modules, specialized for processing specific sensory and modulatory inputs, motor outputs, or sensorimotor integration. This toolkit-like, modular design has been already adopted in some studies of larva behavioral neuroscience because it offers a high degree of configurability, enabling researchers to compare models by adding, removing, or replacing modules [7]. It also facilitates expansion through the seamless integration of new modules.

The hybrid nature of the framework imposes minimal constraints on the modeling detail within individual modules. Whether deterministic, stochastic, rule-based, rate-coded neural models, or neuron-level spiking models, modules can be combined, replaced, and compared, provided they conform to standardized input-output formats. Once a modular larva model has been assembled, it can undergo a genetic algorithm optimization process prior to its use in evaluation studies.

The following sections elaborate on the implementation, usage, and accessibility of the platform in greater detail. Section Design and Implementation describes the architecture and implementation of the Larvaworld platform, including its modeling principles, simulation modes, configuration options, and analysis tools. Section Results presents a range of scientific applications, demonstrating how Larvaworld has been used for behavioral modeling, data-driven analysis, and model evaluation in various settings. Section Code availability provides information on code availability and installation. A schematic of the main components is shown in Figure 1 and a summary of preconfigured virtual experiments is included in Table 2, offering quick-start examples for further exploration.

**Fig 1.**
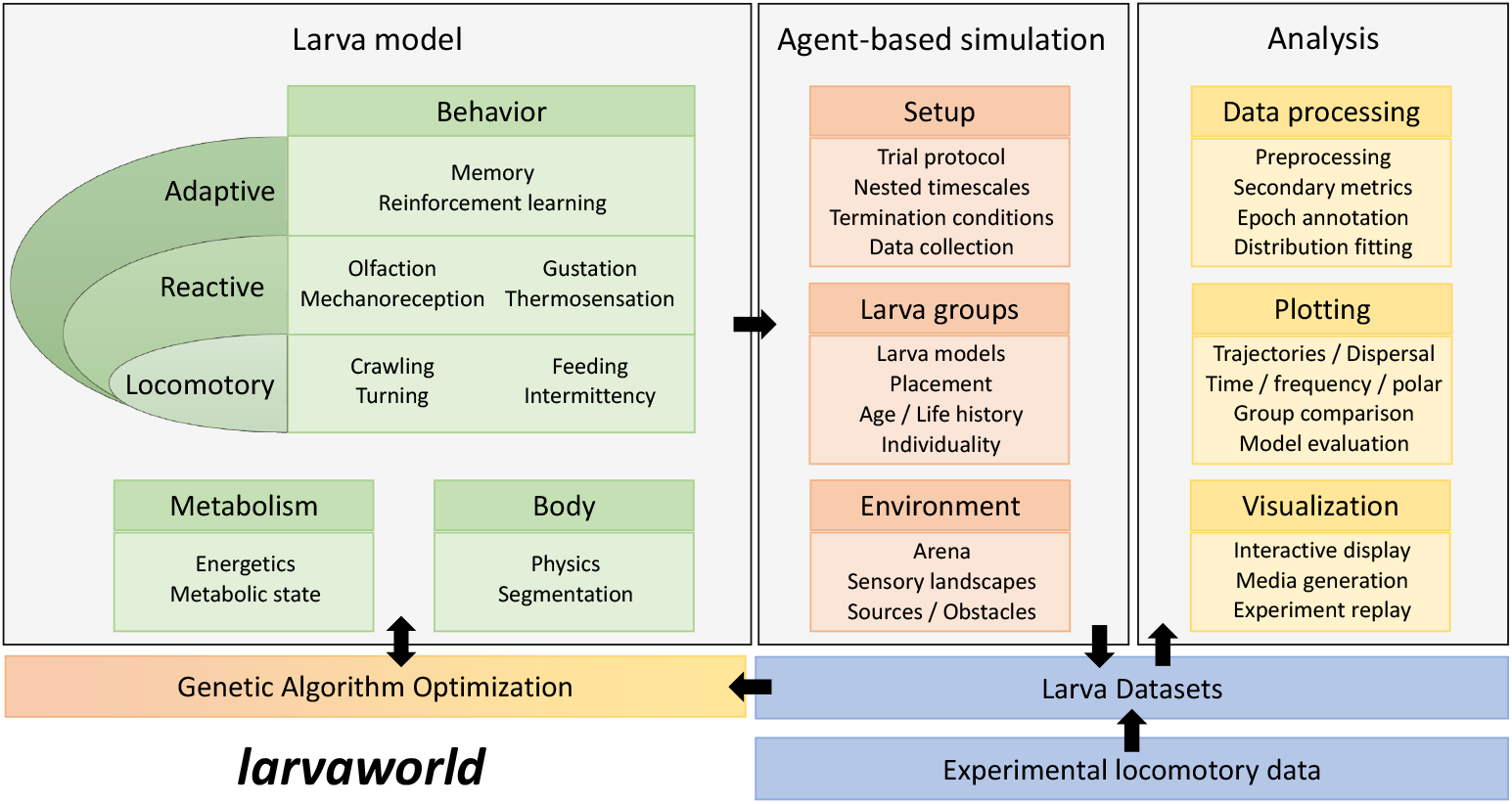
*Larvaworld* architecture. A schematic of the main components and functionalities of *Larvaworld*.

## Design and Implementation

### Modeling principles

The design of the Larvaworld platform addresses four aims:

- Integration of established theoretical and modeling principles from distinct fields across behavioral sciences
- User-friendly interface for both behavioral modeling and analysis purposes
- Modularity and extendability
- Computational efficiency and storage management

In the following sections the motivation behind each developer decision will be articulated along with the specific implementation choices.

### Agent-based modeling (ABM)

The platform is designed to integrate established computational paradigms from diverse research fields, with a primary focus on the agent-based modeling (ABM) approach, which has been extensively used in computational ecology [8, 9]. This choice is motivated by theoretical and practical considerations. ABM provides a powerful framework for simulating complex, dynamic systems where individual agents -representing organisms, behavior, or components- interact with each other and with their environment, making it particularly well-suited for modeling the behavior of *Drosophila* larvae in a virtual space.

The core simulation and agent classes in *Larvaworld* are built upon the *agentpy* package, a Python-based ABM framework and a key dependency of the platform [10]. *Agentpy* ‘s basic *Model, Space* and *Object* classes are adjusted to meet the specific needs of *Larvaworld* especially the nested-dictionary implementation of agent parametrization, ensuring that the platform can accommodate the unique requirements of simulating complex modular biological systems. ABM provides a flexible and efficient structure for the creation and management of agents, including the ordering of their actions, simulation setup, and workflow, and turn-based data retrieval.

### The Dynamic Energy Budget (DEB) theory

To bridge the gap between the neuroscientific and homeostatic timescales of behavioral modeling, *Larvaworld* can place fast neural control of behavior under the regulation of a slower energetics model which addresses energy allocation according to the metabolic needs of the virtual animal as these develop during the larval life stage. The energetics simulation is formulated as a dynamic energy budget (DEB) model following the established literature [11].

The DEB model takes into account the post-hatch age of the larva and the rearing conditions including nutritious substates and starvation or partial food-deprivation periods. It enables the introduction of discrete foraging phenotypes, such as rovers and sitters [12], by differentially configuring the nutrient absorption rate during feeding, eventually regulating the behaviorally expressed exploration-exploitation balance via a dynamic energy-reserve dependent hunger drive. The energetics model runs in the background of the behavioral simulation at an adjusted circadian timescale achieving realistic growth curves for virtual larvae.

### Parametrization

All classes in *Larvaworld* are parameterized using the *param* Python library, a member of the *holoviz* ecosystem, developed to enhance transparent subclassing, interactive visualization through dynamic widgets (see Fig 4), and intuitive real-time display of their parameters within Jupyter notebooks, as demonstrated in the tutorials available on the documentation page (see Figures 6, 7, 5).

These interactive features are illustrated in a number of browser-based applications (see Fig 8) that can later be integrated into a comprehensive Graphical User Interface (GUI) to enhance the accessibility of the virtual laboratory, making it easier to use for researchers and educators without extensive coding experience.

### Visualization

*Larvaworld* uses the *pygame* Python library to support visualization of behavioral simulations as well as replays of previously stored simulations and real-world experiments reconstructed from imported data. These can be run at real-time, slower or faster up to the device’s computational limits. If visualization is enabled for a simulation, a pop-up screen shows the larvae and the environment objects (odor/food sources, impassable borders) on a 2D arena with a realistic spatial scale and timer. Keyboard shortcuts and mouse clicks allow real-time interaction with the visualization screen. Zooming in and out is allowed as well as selecting and locking the screen on specific individuals. Larva unique IDs as well as the screen scale and timer can be toggled on and off. Additionally, larva midline and contour can be toggled, and their head or centroid can be pointed out. Larva trajectories can be shown and their duration can be adjusted. Larvae can be colored with default or random colors or dynamically according to their instantaneous behavior. The arena color can be changed and the distribution of specific odors (odorscape) dynamically visualized. Larvae, sources and borders can be added or deleted instantly. Finally videos of the simulations and replays can be stored, arena or odorscape snapshots captured and videos collapsed on a single image file comprised of all video frames overlaid. A number of snapshots illustrating the real-time visualization of both reconstructed experiments and simulations are shown in Fig 2. A summary of the available online controls is shown in Table 1.

**Fig 2.**
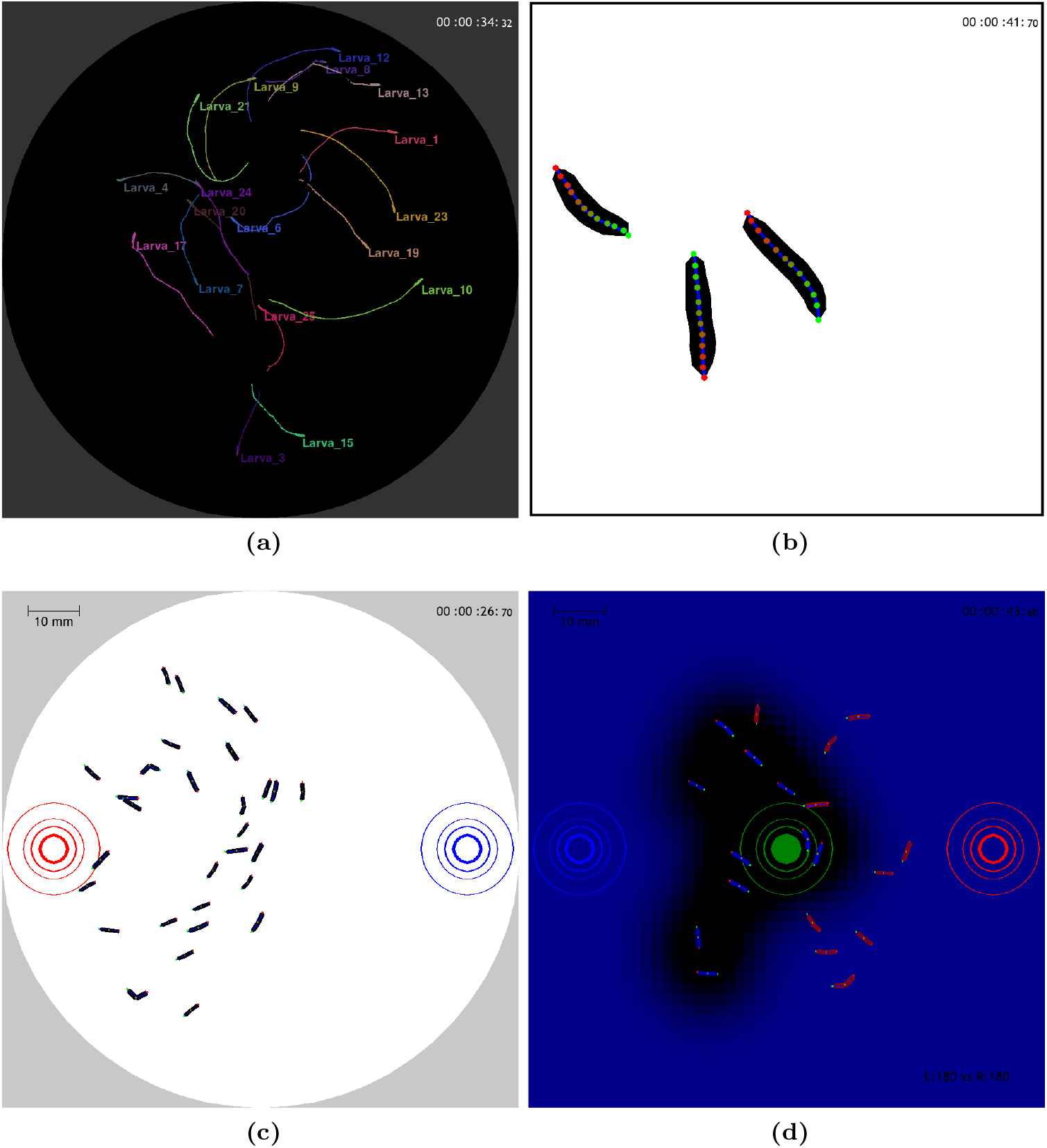
Real-time visualization of reconstructed real-animal experiments and agent simulations. Snapshots from the experimental arena illustrate: (a) real-animal behavior displayed on a black background with visible larval IDs and reconstructed locomotion trajectories [13]; (b) a close-up showing the 12-point midline tracking of individual larvae [13]; (c) a simulated odor preference experiment: A group of larvae is randomly placed at the centre of the dish with an appetitive odor source placed on the left (red) and a non-valenced odor on the right (blue). The larvae are gradually attracted to the side of the appetitive odor; and (d) a capture-the-flag mini-game: Two larva groups (blue on the left VS red on the right) compete to capture a highly-valenced centrally-placed odor source (green) and carry it back to their base (low-valenced odor source of the respective color). The simulated olfactory sensory landscape (“odorscape”) emanating from the green odor source is shown in black over a blue background.

**Table 1.** Visualization default keyboard/mouse controls. L*, R*, M* stand for mouse left, right, and center buttons respectively.

| Screen |  | Drawing |  | Color |  | Interaction |  | Simulation/Storage |  |
| --- | --- | --- | --- | --- | --- | --- | --- | --- | --- |
| Scale | s | midline | m | random | r | select | L* | snapshot | i |
| Timer | t | contour | c | behavior | b | lock<br>screen | f | odorscape | o |
| IDs | TAB | head | h | background | g | delete | del | pause | p |
| Zoom in/out | M* | centroid | e | odorscape | 0-9 | add | L* |  |  |
| Screen Motion | ↕↔ | trail | p |  |  | inspect | R* |  |  |
|  |  | trail<br>duration | +/- |  |  | dynamic<br>graph | g |  |  |

Replays of simulated or imported experimental datasets allow additional configuration. Inclusion of specific individuals and time range can be defined. Replays can be run on arenas different than their original one as long as all tracks remain within their boundaries. To this end, larva x-y coordinates can be transposed to the arena center. Additionally all tracks can be aligned to start from the same origin at the arena center, a mode favoring inspection of larva dispersion over time. Larva trails can be colored according to the instantaneous forward or reorientation angular velocity. For closely inspecting single-larva replays, the screen center can be locked to a specific midline point or segment and a background hue can be enabled to easily assess body movement in space. Finally when replaying imported experiments, larva bodies can be reconstructed as segmented virtual bodies of a given number of segments in order to make them visually comparable to virtual larvae.

### Simulation modes

Several simulation modes are available, each serving a distinct purpose and featuring its own set of configurable parameters. Such a mode-specific parameter set fully configures a simulation and can be stored as a nested Python dictionary under a unique ID for future usage. A rich repertoire of preconfigured parameter sets is already available upon installation consisting of several simulation examples for each mode.

The Command Line Interface (CLI) can be summoned directly from the terminal using the larvaworld command. This has to be followed by an essential argument that defines the desired simulation mode by its respective shortcut. In some modes, a second argument is needed to retrieve an existing, stored configuration by its unique ID. Additional arguments can be used to overwrite some basic parameters of the predefined configuration. The available simulation modes are shown in Fig 3 while their usage via CLI commands is showcased separately.

**Fig 3.**
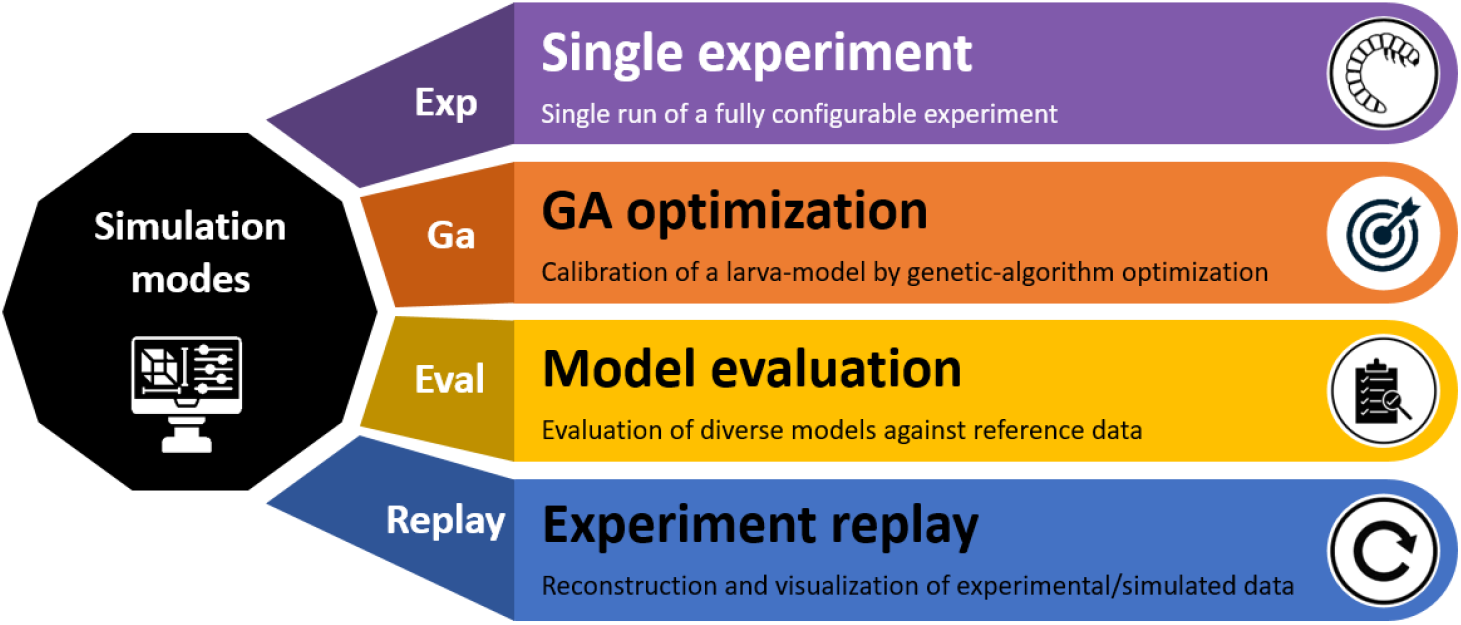
CLI simulation modes. The simulation modes available in *Larvaworld* along with the respective argument to launch them via the command-line interface.

### Single experiment

This is the basic simulation mode which runs a single virtual experiment once, based on a detailed configuration of the virtual environment, involved larva groups, temporal simulation features and subsequent analysis pipeline. Such a configuration might aim to enable the detailed, standardized, realistic, in silico replication of a real-world experiment, in which case its setup and analysis is drawn from published behavioral studies or established behavioral protocols. Such examples can be found among the multiple preconfigured experiments listed below (Table 2). Alternatively, it might define a difficult-to-implement or entirely fictitious experiment for model-testing, visualization or proof-of-concept purposes. As a distinct simulation mode, an experiment launched individually supports a complete pipeline to analyze the results, generate the respective plots and store the simulated datasets for future access. In contrast, the single run forms the core simulation unit, the building block for more complex simulation modes described below.

**Table 2.** Summary of preconfigured behavioral experiments.

| Type/Behavior | Experiment | Description | Literature source |
| --- | --- | --- | --- |
| exploration | close-view | Single larva closely inspected in a tiny arena | - |
|  | dish | Exploration of a non-nutritious Petri-dish | - |
|  | dispersion | Larva dispersion from the arena center | - |
| chemotaxis | navigation | Navigation up an odor gradient | Gomez-Marin et al. (2012) [14] |
|  | local search | Exploration in the vicinity of an odor source | Gomez-Marin et al. (2012) [14] |
| odor preference | train & test | Olfactory associative learning (train & test phase) | The Maggot Learning Manual |
|  | test on/off food | Test in the presence/absence of nutritious substrate | The Maggot Learning Manual |
| foraging | patchy food | Foraging in arena with one/two/multiple food patches | - |
|  | uniform food | Foraging in uniformly distributed nutritious substrate | - |
| growth | rearing | Larva rearing in ad-libitum conditions | - |
|  | rovers VS sitters | Foraging phenotypes compared in diverse conditions | Kaun et al. (2007) [12] |
| imitation | realistic bodies | Multisegment larvae in Box2D physics engine | - |
|  | dataset imitation | Experimental dataset imitation | - |
| games | maze | Navigation in a maze towards an odor source | - |
|  | capture/keep the flag | Larva teams competing for a portable nutritious object | - |

Each of these lines runs a dish simulation (30 larvae, 3 minutes) without analysis:

~~~
larvaworld Exp dish −N 30 −duration 3.0 −vis_mode video
larvaworld Exp patch_grid −N 30 −duration 3.0 −vis_mode video
~~~

This line runs a dispersion simulation and compares the results to the existing reference dataset. We choose to only produce a final image of the simulation.

`larvaworld Exp dispersion −N 30 −duration 3.0 −vis_mode image −a`

### Genetic Algorithm for model optimization

As explained above, the *behavioral architecture* applying neural control over an agent’s behavior is hierarchically layered and modular. The latter implies that diverse mutually exclusive models can compete for each module defined within the architecture. The nature of these competing module-specific models is not constrained apart from the need to adhere to the input/output specification of the module. In other words each of these candidate models might entail an intrinsic set of configuration parameters, not shared among other candidate models. An issue that surfaces when comparing across them, is that each model should be represented by the best possible parameter configuration, so that the comparative simulation study indeed selects between optimal model instances. It follows that an optimization process for each candidate model that will yield its optimal representative, must precede the comparative study. This need has motivated the integration of a genetic algorithm (GA) optimization process into *Larvaworld*.

The GA simulation mode accepts three sets of parameters as illustrated in Fig 4 :

**Fig 4.**
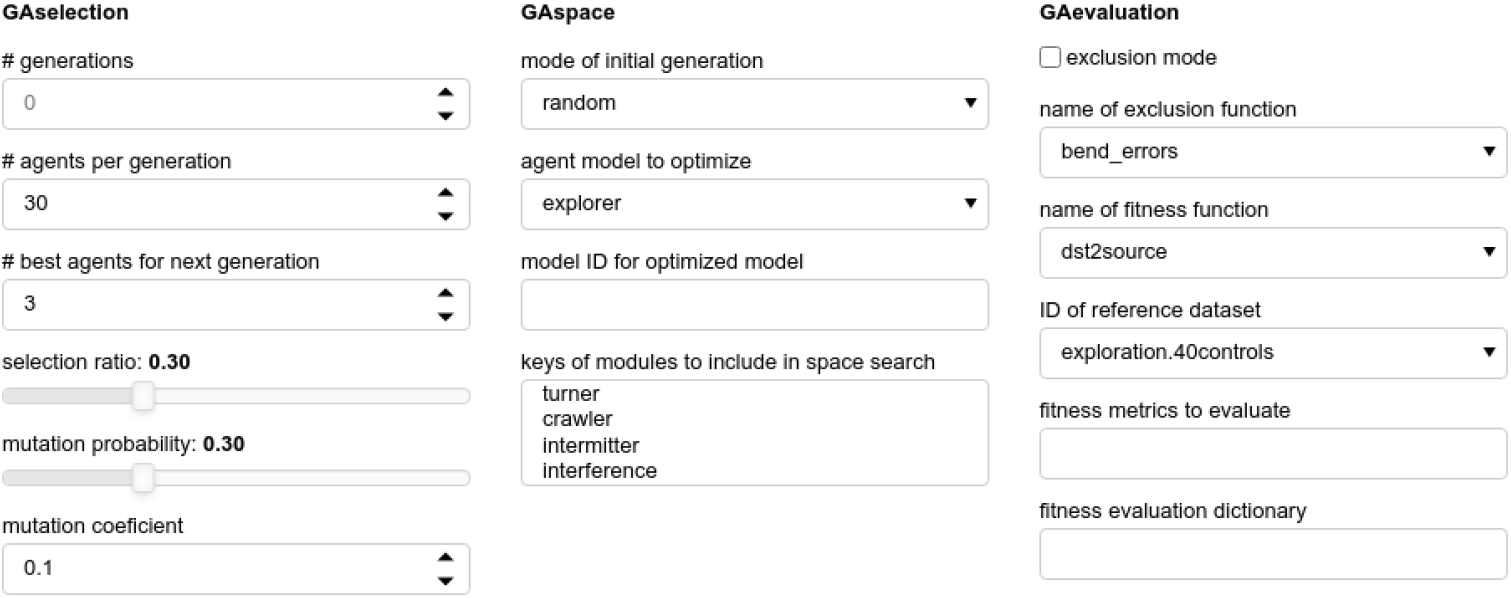
GA configuration panel

- **Selection algorithm** : Here we define the number of generations to run and the number of agents per generation. Additionally, the criteria to select and optionally mutate agent configurations to generate the next generation.
- **Parameter space** : A basic model configuration needs to be provided along with the specification of the parameters that will be optimized. Each of the parameters-to-be optimized is part of an existing module of the agent’s behavioral architecture and the allowed range of the space search is predefined.
- **Performance evaluation** : Parameters that ultimately define a fitness function that quantifies the perfomance of each agent in each generation. In the simplest case the fitness function is defined externally and provided as an argument. Alternatively, it may be constructed by the GA engine based on provided parameters. For example when the agents are evaluated against an experimental reference dataset.

This line optimizes a model for kinematic realism against a reference experimental dataset :

~~~
larvaworld Ga realism −refID exploration.30 controls −N 20 −duration 0.5
    *↪* −mID1 GA_test_loco −init_mode model
~~~

### Model evaluation - comparison with real data

Sometimes we focus on evaluating our agent models against some reference experimental dataset. The model evaluation mode does exactly that. As shown in Fig 5, it requires defining :

**Fig 5.**
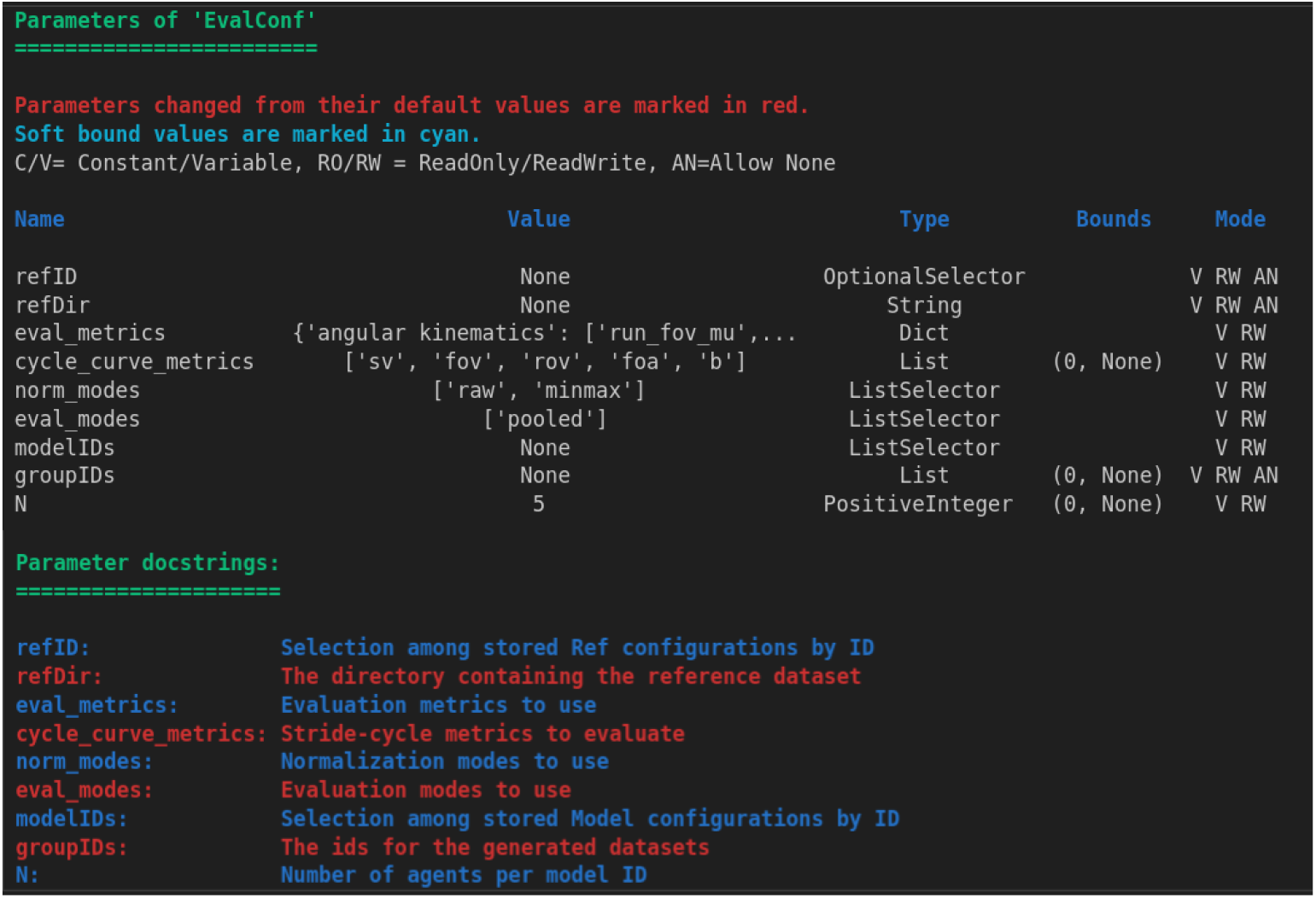
Model Evaluation configuration parameters

- Reference dataset, designated via ID or directory
- Larva models retrieved via ID and the size and IDs of the respective larvagroups
- Evaluation metrics and configuration
- Simulation experiment and its setup

Illustrative examples of model evaluation simulations are presented in the Results section. This mode is available via CLI as well. The following line evaluates two models against a reference experimental dataset:

~~~
larvaworld Eval −refID exploration.30 controls −mIDs RE_NEU_PHI_DEF
    *↪* RE_SIN_PHI_DEF −N 3
~~~

### Experiment replay

Replay a real-world experiment. This line replays a reference experimental dataset :

~~~
larvaworld Replay −refID exploration.30 controls −vis_mode video
larvaworld Replay −refDir Schleyer Group/processed/exploration/30controls
     *↪* −vis_mode video
~~~

## Simulation configuration

### Environment setup

The simulation environment in *Larvaworld* is designed as a spatial arena with configurable shape and dimensions. Within this arena, single or grouped odor/food sources and impassable borders can be placed at specified locations. A key feature of the environment is its customizable sensory landscape, which includes gradients of various sensory modalities such as olfactory (odorscape) emanating from odor sources placed inside the arena, thermal (thermoscape), and wind (windscape). The parameters of the respective class are shown in Fig 6. Mini videos of windscape and odorscape are available online [15–19].

**Fig 6.**
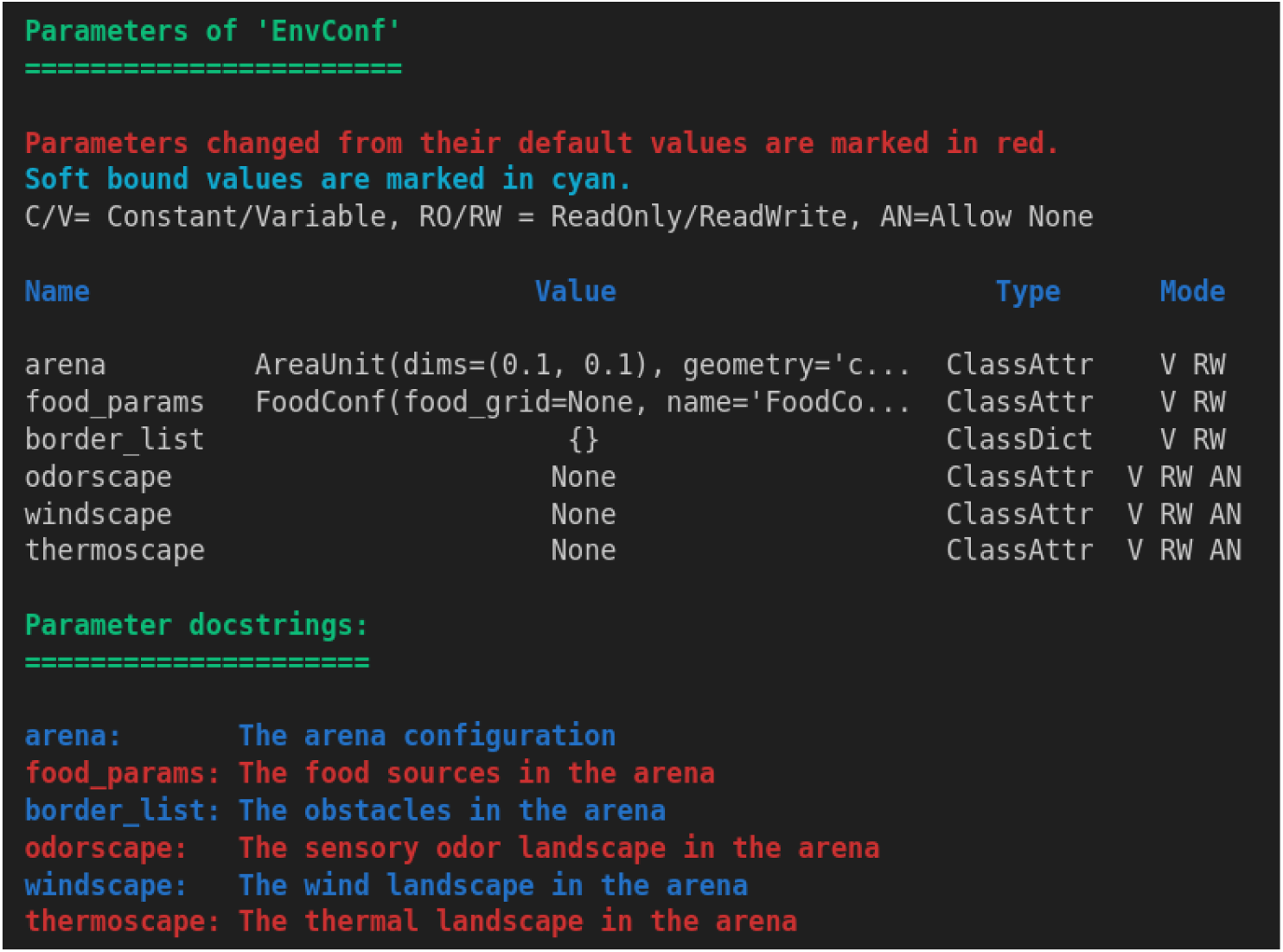
Virtual environment configuration parameters

To facilitate diverse experimental setups, multiple predefined environmental configurations are available, each of which can be adjusted to suit specific research needs.

Researchers can modify attributes such as the position and intensity of sensory sources, the spatial distribution of sensory gradients, and the boundaries of the arena. This flexibility enables precise control over the simulation environment, allowing for the design of experiments that replicate real-world conditions or explore novel scenarios.

### Substrate

Larva rearing and behavioral lab experiments are carried out on diverse nutritious or non-nutritious substrates. Several substrate types have been established in terms of their nutrient compound composition. Some of these have been implemented in *Larvaworld* with their characteristic composition. Virtual larvae can be grown, starved or tested on such substrates (Table 3). Any food source in the arena is characterized by a substrate of specific type, nutritional quality and available amount of food. Alternatively, the substrate can be placed locally as patches in a grid that covers the entire arena as a grid where each cell can independently hold a certain amount of food. This food can be detected and consumed until depleted.

**Table 3.** Compound composition of established nutritious arena substrates.

| Substrate | Compound density ($\mu g/ml$) | | | | | | Literature source |
| --- | --- | --- | --- | --- | --- | --- | --- |
|  | glucose | dextrose | saccharose | yeast | agar | cornmeal |  |
| standard-medium | 100 | - | - | 50 | 16 | - | Kaun et al. (2007) [12] |
| PED-tracker | - | - | 10 | 187.5 | 5000 | - | Schumann et al (2020) [20] |
| cornmeal | - | 70.3 | - | 14.1 | 6.6 | 65.6 | Wosniack et al. (2021) [21] |
| sucrose | 17.1 | - | - | - | 4 | - | Wosniack et al. (2021) [21] |

### Larva groups

In *Larvaworld*, virtual larvae are generated in groups, with members of each group sharing specific traits that distinguish them from individuals in other groups. These traits include not only the configuration of the larva model itself but also elements such as a shared life history, a spatial distribution that defines their initial pose (Table 4), a group-specific color for easy visualization and a distinct odor signature (Fig 7).

**Table 4.** Larva group initial spatial placement parameters.

| Parameter | Description |
| --- | --- |
| N | Number of virtual larvae in the group |
| location | Centre of the spatial distribution in the arena |
| scale | Spatial extent of the distribution |
| shape | Shape of the distribution (circular/rectangular/oval) |
| placement | Placement within the distribution’s shape (uniform/normal/periphery) |
| orientation | Range of initial spatial body orientations |

**Fig 7.**
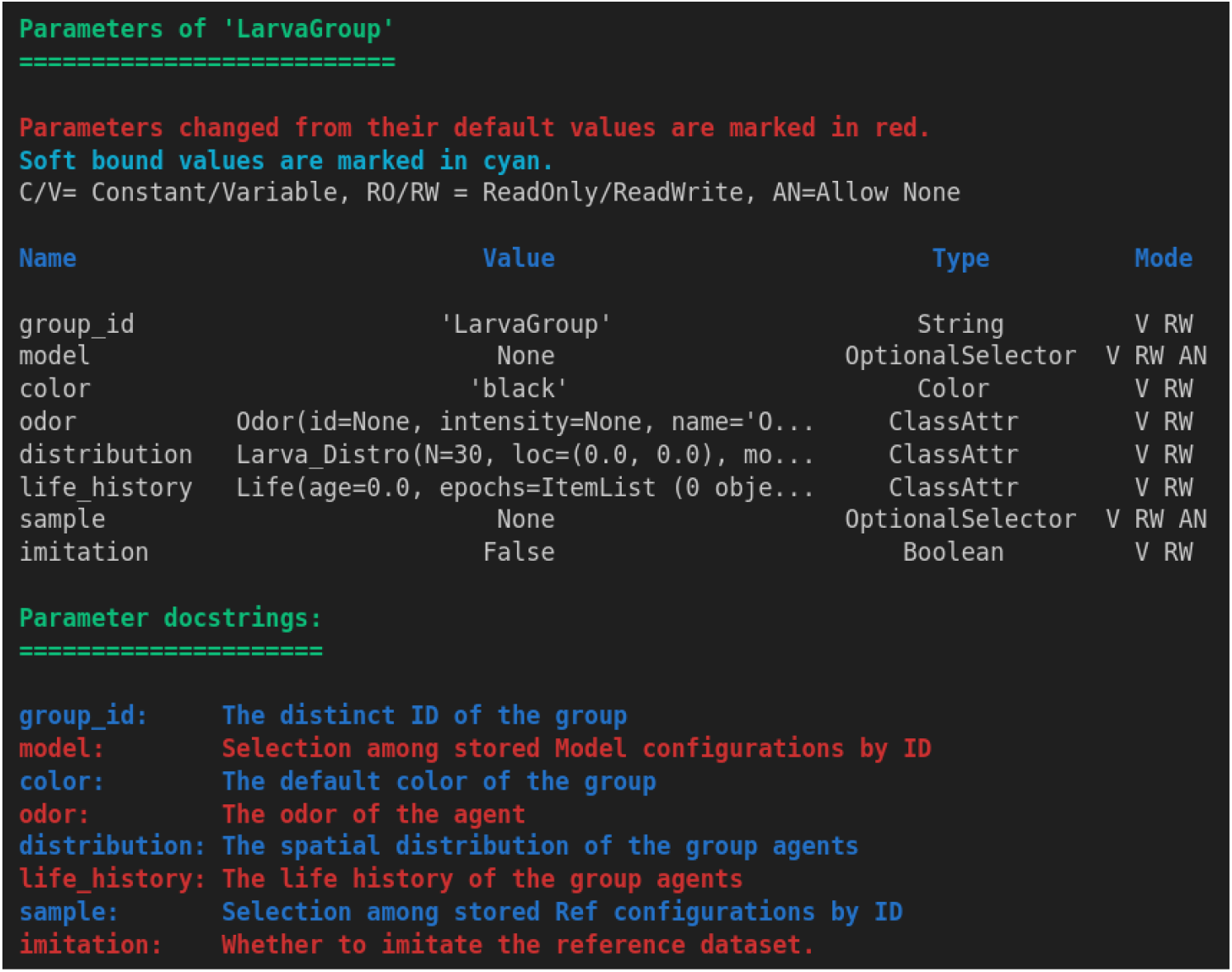
Virtual larva group parameters

Virtual larvae can have their own life history at the moment they enter the experimental arena. To simulate that, a *Drosophila*-specific dynamic energy budget (DEB) model simulates larva growth up to a specific age post-hatch, on a predefined rearing substrate (Table 3) of specified quality so that the behavioral simulation can be subsequently initiated. Periods of food deprivation or complete starvation during rearing or during the actual behavioral simulation can also be defined.

Groups can be assigned a linked reference dataset from which parameters are sampled. The sampling mode is highly configurable, offering three main options: optimizing for an average individual, preserving interindividual variability, or replicating the reference dataset on an individual-by-individual basis for specified parameters. For a detailed comparative evaluation of these group-generation modes, see *Individuality and Variability* in the *Results* section.

This functionality provides researchers with the flexibility to design experiments featuring multiple, distinguishable groups of virtual larvae. By enabling group-specific traits and parameter configurations, *Larvaworld* supports a broad spectrum of experimental scenarios, ranging from comparative analyses to simulations that emulate the diversity and dynamics observed in real-world populations.

## Web-based applications

A number of web-based *Larvaworld* applications are available to facilitate inspection, configuration and real-time visualization of various elements. The panel showing the available applications can be launched via the CLI command `larvaworld-app`. Some of the available applications are listed in Table 5 and a snapshot is shown in Fig 8.

**Table 5.** Web-based applications.

| Application | Description |
| --- | --- |
| Experiment Viewer | Inspect/launch preconfigured experiments |
| Larva Models | Inspect/visualize modular larva-models |
| Locomotory Modules | Inspect/test behavioral modules |
| Track Viewer | Visualize stored datasets |

**Fig 8.**
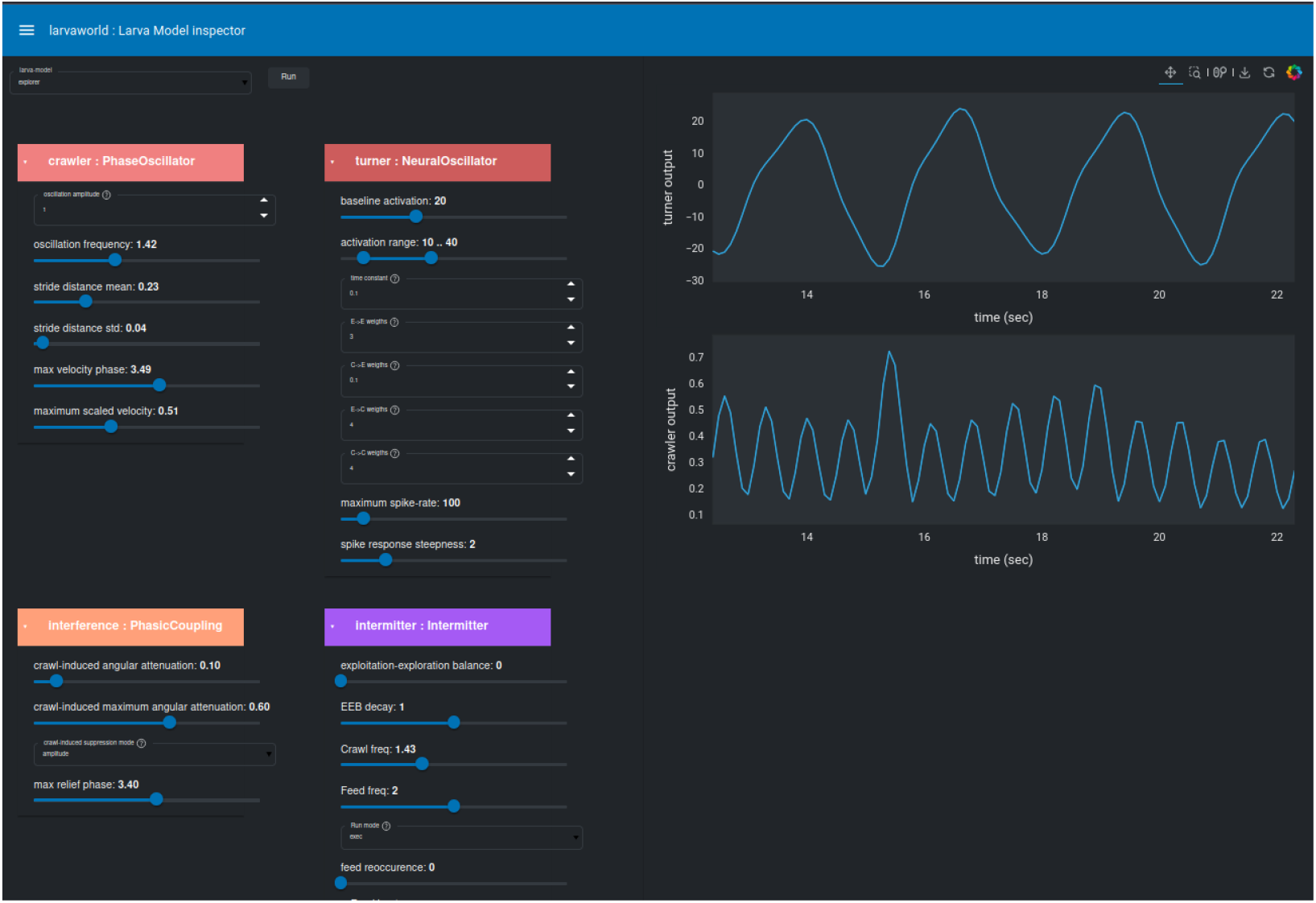
Web-based *Larvaworld* application to inspect the modular composition of any preconfigured locomotory larva-model. This can be selected by its unique ID from a drop-down list of all available models. The configuration of all 4 basic modules comprising the locomotory layer of the model is available on the left sidebar. Real-time simulation variables (e.g. input/output) are dynamically plotted in the middle.

## Data management and analysis

### Standardized format for both experimental and simulated datasets

*Larvaworld* is set up primarily to analyze and simulate *Drosophila* larva motion-tracking experiments. In such studies recorded data are usually collected per trial and per animal group. Datasets are then comprised of a number of (identical) trials for each of a number of animal groups (for example different genotypes, satiation states, rearing conditions) and their control counterparts. Furthermore it is common to perform the same behavioral experiment under a number of different environmental conditions or even to compile an essay of diverse behavioral experiments across animal groups. To make the importation of such experimental datasets straightforward to the user, the platform’s data organization builds on such a *LarvaDataset* as a core class. Crucially this class is common for both simulated and experimental data in order to facilitate comparison and model fitting. The basic elements of a *LarvaDataset* instance are :

- Timeseries data computed at every timestep for every larva are stored as a double-indexed *Pandas* dataframe where the index defines the timestep and the unique ID of the larva, while the values of each parameter are stored in a separate column. Initially the dataframe contains only the primary tracked parameters (x-y coordinates of at least the centroid, often a number of points along the larva midline and optionally the contour of the larva body). The dataframe is enriched with derived parameters during data-processing as described in the section below.
- Endpoint metrics computed once for every larva at the end of the simulation are also stored as a *Pandas* dataframe indexed by the agent’s unique ID. This is also enriched during data-processing with summary measurements of the timeseries data.
- Metadata about experimental conditions, tracking parameters, animal groups etc are structured as a nested dictionary. Storage paths are also stored there for easy access during analysis.

To facilitate efficient data storage, retrieval, and analysis the dataframes are stored in an *HDF* file under different keys (e.g. midline, contour, trajectory) for easy access. The metadata dictionary is stored as a configuration text file alongside the data. The dataset can optionally be registered as a reference dataset, by providing a unique ID as key, a functionality often used to easily access real experimental data during model evaluation and optimization but also for quickly visualizing comparative plots of simulated and experimental data. Additional files are created during data-processing.

### Unbiased parameter computation and data analysis from primary 2D timeseries

In *Larvaworld*, data processing methods are identical for both simulated and imported experimental data, ensuring consistency and unbiased analysis. The platform supports three sequential, configurable pipelines that can be applied to any dataset containing at least the x-y coordinates of a single point per larva (Table 6):

**Table 6.**
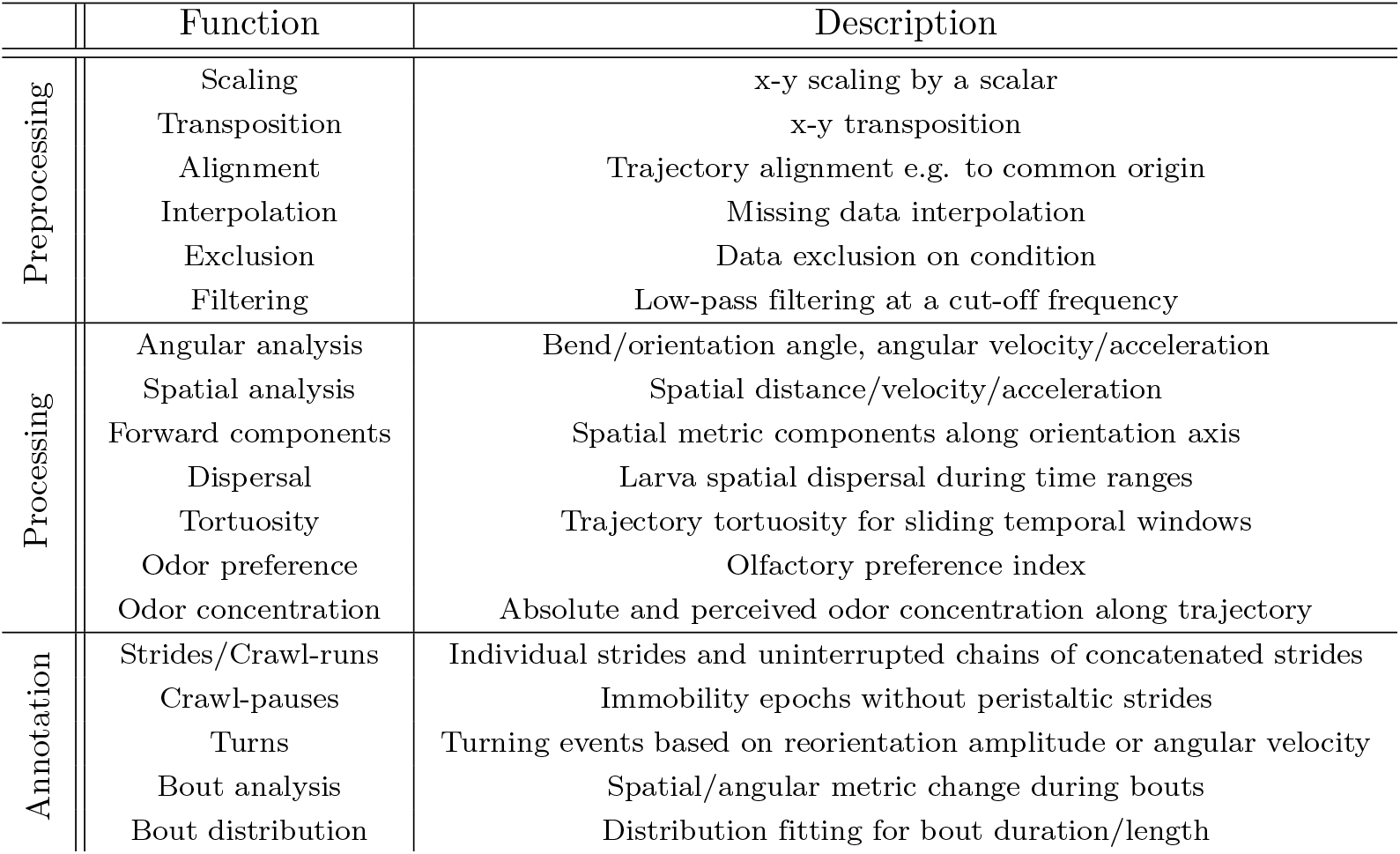
Data processing methods.

During preprocessing the dataset x-y series can be rescaled by a scalar (e.g. to convert datasets stored in millimeters to the default *Larvaworld* space unit of meters) or transposed to align with the arena center. Datasets can also be aligned to a common origin, facilitating visualization of dispersal patterns. Noise from tracking artifacts can be reduced using low-pass filtering at a configurable cut-off frequency. Additionally, missing data points can be interpolated, or specific data slices can be excluded based on predefined conditions (e.g. instances of larva collisions).

In the main processing stage, secondary parameters are derived from the primary x-y time series:

- Angular analysis: Bending angles and absolute orientation in 2D space are computed for individual larval segments. Instantaneous orientation and bend can be defined either directly from these angles or via front/rear body vectors.
- Spatial analysis: Key metrics such as distance, velocity, and acceleration (including their components along the larva’s forward orientation axis) are calculated.
- Dispersal patterns over specified time intervals
- Trajectory tortuosity using sliding windows of configurable durations
- Odorscape Navigation: Metrics such as instantaneous odor concentration and perceived concentration changes along the larva’s trajectory are available. Additionally it is possible to track the instantaneous distance and bearing to odor or food sources.
- Preference index calculations for olfactory preference experiments.

Finally the bout annotation pipeline identifies discrete behavioral events, including:

- Strides: Individual locomotor steps.
- Crawl-Runs: Continuous sequences of strides.
- Crawl-Pauses: Periods of immobility.
- Turns: Defined either by reorientation amplitude or changes in orientation/bending angular velocity sign.

A fitting algorithm determines optimal distributions for the temporal durations or spatial lengths of bouts, selecting from options such as power-law, exponential, and log-normal distributions. Additional metrics include distance, orientation change, and bending angle change during individual bouts, as well as crawling frequency estimates. For virtual larvae, the platform enables tracking of a broad range of model-specific, simulation-derived metrics, offering flexibility for detailed behavioral analysis and model evaluation.

### Importing experimental datasets from diverse tracker setups

Experimental laboratories employ a wide range of tracking software to accurately capture 2D larval locomotion. Typically, primary data consist of coordinates for multiple points along the body midline and around the body contour, tracked at either fixed or variable frame rates. These raw data are exported in tracker-specific formats and subsequently processed to derive secondary parameters for analysis. *Larvaworld* supports the direct import of tracked datasets from a variety of tracker-specific formats (Table 7). For each supported format, the platform defines key parameters such as the tracking frame rate and the number of midline or contour points, along with a format-specific conversion function to transform the raw data into the standardized LarvaDataset format. Users can specify which tracks to include using a set of import arguments, such as track duration, start or termination time, and the option to limit the dataset size by defining a maximum number of animals. The imported datasets are standardized, making them directly comparable both to each other and to simulated data. Importantly, to promote reproducibility, only primary parameters are imported, ensuring that all subsequent derived metrics are transparently calculated within the platform. This approach facilitates consistent and unbiased comparisons across experimental and simulated datasets while maintaining a high degree of accessibility and flexibility for data analysis.

**Table 7.**
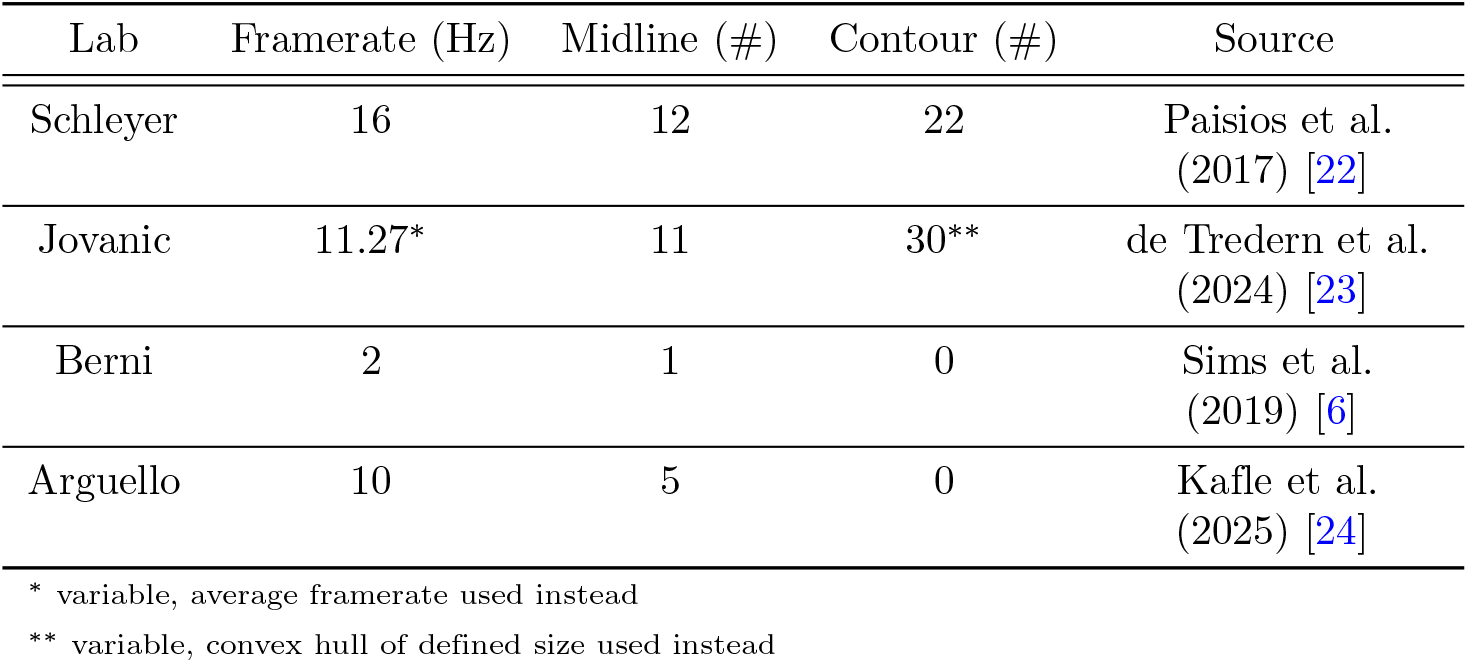
Lab-specific experimental data-formats.

## Results

### Scientific applications

The *Larvaworld* package has already been successfully used in several scientific studies. These will be briefly described here, focusing on the way each of them used *Larvaworld*.

As a whole, these studies illustrate the spectrum of functionalities supported by the package, both as a data analysis and a behavioral modeling tool.

The simplest and most straightforward case is usage of Larvaworld as a data analysis tool. A study of feeding-state dependent aversive behavioral responses of *Drosophila* larvae to mechanosensory stimulation aimed to elucidate neuropeptidergic modulation of reciprocally interconnected inhibitory neurons [23]. The tracker-specific datasets of recorded larva locomotion across all experimental groups were successfully imported into *Larvaworld*. Using the platform’s comparative analysis methods, differences in several key kinematic locomotory parameters were identified among fed, starved, and sucrose only fed groups. The analysis was further extended to examine locomotory differences in dehydrated (sucrose only fed) and rehydrated, as well as food-deprived and refed animals, across multiple genetically distinct strains. This comprehensive approach allowed for a detailed assessment of how nutritional and hydration states influence locomotory behavior.

The *behavioral architecture* framework that forms the backbone of the modular modeling functionality in Larvaworld has been presented in [4]. In this study both data analysis and modeling is done via *Larvaworld*. Novel insights derived from the analysis of experimental datasets are utilized in order to extend a previously published locomotory model of the larva [25] with two novel features, namely crawl-bend interference and behavioral intermittency, the latter also based on a previous modeling study [26]. The resulting intermittent coupled-oscillator model of larva locomotion is the default model in *Larvaworld* for performing behavioral simulations in stimulus free or sensory-rich environments, in the latter case augmenting the locomotory model with sensors to detect e.g. odor or food sources.

The modular approach to behavioral modeling featured in *Larvaworld* is showcased in a study of associative learning and conditioned behavior of the *Drosophila* larva [27]. The study features a spiking model of the larva’s mushroom body (MB), a neuropile that has been implicated in memory and learning. In order to evaluate the MB model in realistic odor-preference simulations, and to compare the results to lab experiments, it has been plugged in as a memory module. This highlights how the modular architecture of *Larvaworld* contributes to answering relevant research questions across domains.

Notably the resulting modular larva model was a hybrid also in the sense that its comprising modules operated in nested timescales, namely the spiking MB module at a 0.1 msec timestep whereas the behavioral locomotory simulation at the default 0.1 sec timestep. This is an illustration of the capability of the behavioral architecture to integrate disparate computational modules regardless of their level of detail or temporal resolution.

Moreover the modular modeling approach facilitates extendability. In a comparative evolutionary study of thermotaxis across drosophilids, larvae from 8 distinct species have been tested in a suitably configured thermal-gradient arena (thermoscape) in order to quantify and compare their preferred temperatures [24]. Simulations were used to evaluate competing hypotheses about the neural underpinnings of the observed temperature preferences across species. In order to couple real animals to their virtual counterparts, species-specific models were fitted. A virtual thermoscape arena was configured and the default locomotory model was extended with an additional thermosensation modality. The performed temperature-preference simulations closely resembled the experiments in their spatiotemporal configuration, larva groups and behavioral results, while additionally allowing the evaluation of competing circuitries of thermotaxis models.

### Import and analysis of experimental datasets

This example illustrates the process of importing datasets of *Drosophila* larva locomotion. The goal is to compare the locomotion patterns of larvae subjected to different diets over 5 hours : normally fed, fed only with sucrose (therefore protein-deprived), and completely starved. The distinct metabolic states of the larva groups might have an impact on their locomotion. We specifically focus on the temporal evolution of their dispersal in space.

The import process involves converting the raw tracker-specific data format into the *Larvaworld* format. This conversion is performed only once, and the processed data is stored for future use without the need for repeated conversions.

Once the datasets are imported, various visualizations can be generated to analyze the locomotion patterns of the larvae. The trajectories of the larvae are plotted, showing their movement over time. Boxplots of endpoint metrics such as mean, final, and maximum dispersal during the first 60 seconds are generated to provide a statistical comparison of the groups. In addition, the dispersal of larvae from their starting point is plotted over time, capturing the mean and variance of their movement (see Fig 9). Replay simulations are run to visualize the trajectories of the larvae aligned at the origin, and videos of these simulations are generated and combined into a single video.

**Fig 9.**
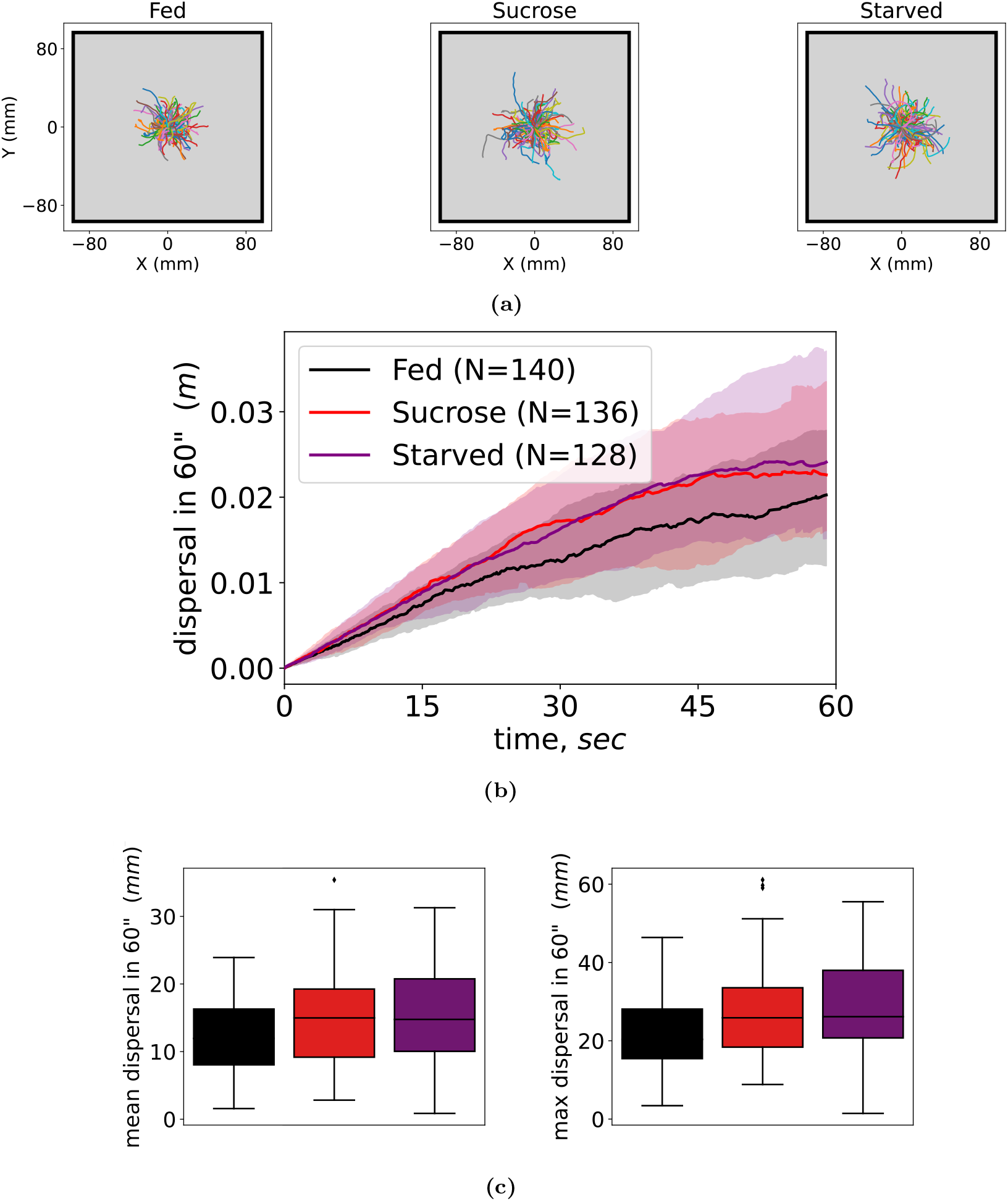
Impact of metabolic state on spatial dispersal analyzed in lab experiments. Three larva groups subjected to different dietary conditions are compared in terms of their spatial dispersal in a quadratic arena during the first minute of the experiment: (a) The trajectories of the larvae, aligned to start from the center of the arena. (b) Temporal course of the larva spatial dispersal. Line indicates the group median while shaded area denotes first and third quartiles. (c) Boxplots of average and maximum dispersal. **Video:** The temporal course of dispersal for the three groups shown in (a) [28].

The combined video output from the replay simulations provides a visual comparison of the locomotion patterns of the three larva groups, showcasing the impact of different metabolic states on larval locomotion. The complete tutorial notebook, including the code and detailed steps, is available in the software’s documentation.

### Demonstration of a chemotaxis essay

To illustrate the usability of the virtual lab, we present a demonstration of a chemotaxis assay, replicating two well-established experimental paradigms [14]. In the first experiment, both the odor source and the larvae are positioned at the center of the arena, where the larvae are expected to exhibit zigzagging or orbiting trajectories around the source, remaining in its vicinity. In the second experiment, the larvae and the odor source are placed on opposite sides of the arena, prompting the larvae to approach the source.

We focus on two distinct chemotaxis algorithms that have been proposed in the literature. The first, as described in [25], suggests that lateral bending is driven by continuous oscillations, which are dynamically modulated by incoming olfactory stimuli. This model lacks behavioral intermittency, resulting in uninterrupted crawling along curved trajectories. In contrast, an alternative model proposes that larvae interrupt their crawling when navigating down an odor gradient to reorient themselves toward the appetitive odor source [29]. This finding can be integrated as an extension of the Lévy walk model, adjusting turning probabilities based on incoming sensory stimulation, as acknowledged in previous studies [30]. The model configurations are summarized in Tab 8.

**Table 8.** Chemotaxis essay. Five locomotory models are tested in a chemotaxis essay. The two basic models employ different algorithms for turning behavior, namely “Tastekin” is a stochastic Levy-walker with the ability to interrupt locomotion when navigating down-gradient, while “Wystrach” features a lateral oscillator generating continuous lateral bending, modulated by the incoming olfactory stimulation. The models are tested using either a logarithmic or a linear function of instantaneous concentration change for sensory transduction. An olfactory-blind model is used as control.

| Model configurations | Tastekin<br>(log) | Tastekin<br>(lin) | Wystrach<br>(log) | Wystrach<br>(lin) | controls |
| --- | --- | --- | --- | --- | --- |
| olfaction sensory<br>transduction mode | logarithmic | linear | logarithmic | linear | logarithmic |
| ability to interrupt<br>locomotion | True | True | False | False | False |
| appetitive odor<br>valence | True | True | True | True | False |

In *Larvaworld* simulations, multiple models can be evaluated simultaneously, with each model represented by a dedicated group of virtual larvae. Utilizing this functionality, we implemented two variations of each chemotaxis model by modifying the sensory transduction algorithm, which translates changes in odor concentration into olfactory stimulation. Additionally, we included an odor-blind control group to serve as a baseline for comparison.

The results of the chemotaxis assay are presented in Fig 10-11. Analysis of the simulated trajectories and corresponding absolute concentration time plots indicates that the intermittent Lévy walk model demonstrates superior performance in both experiments. Meanwhile, the lateral oscillator model performs better than chance in the approach experiment. It is important to note that this demonstration does not incorporate any model optimization step (see *Genetic Algorithm Optimization* in *Simulation Modes*), and thus, further parameter adjustments could potentially improve model performance.

**Fig 10.**
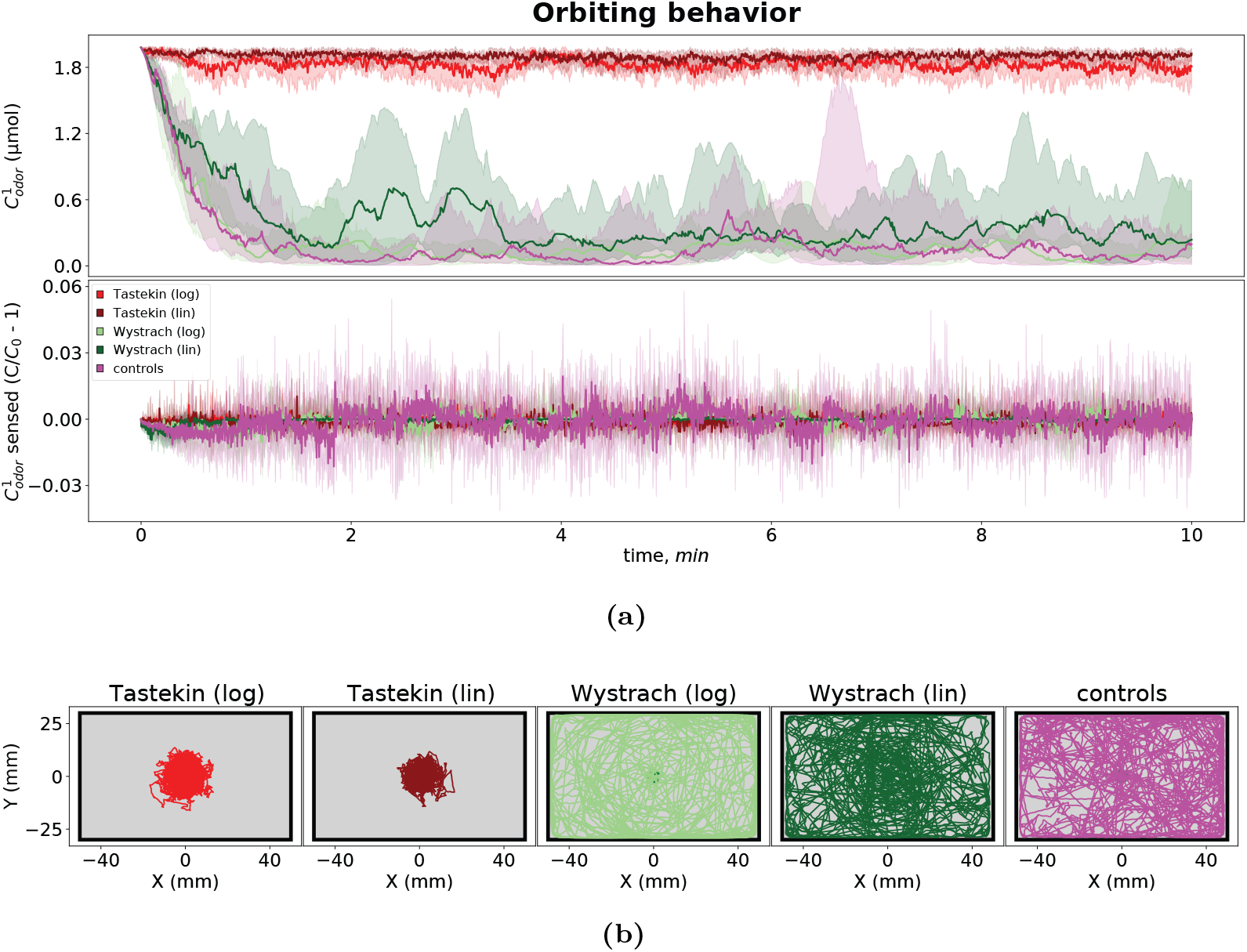
Chemotaxis around the odor source. (a) Time plots of the absolute odor concentration (top) and perceived odor concentration change (bottom) encountered by larvae during the experiments, along with (b) the larva trajectories around the odor source.

**Fig 11.**
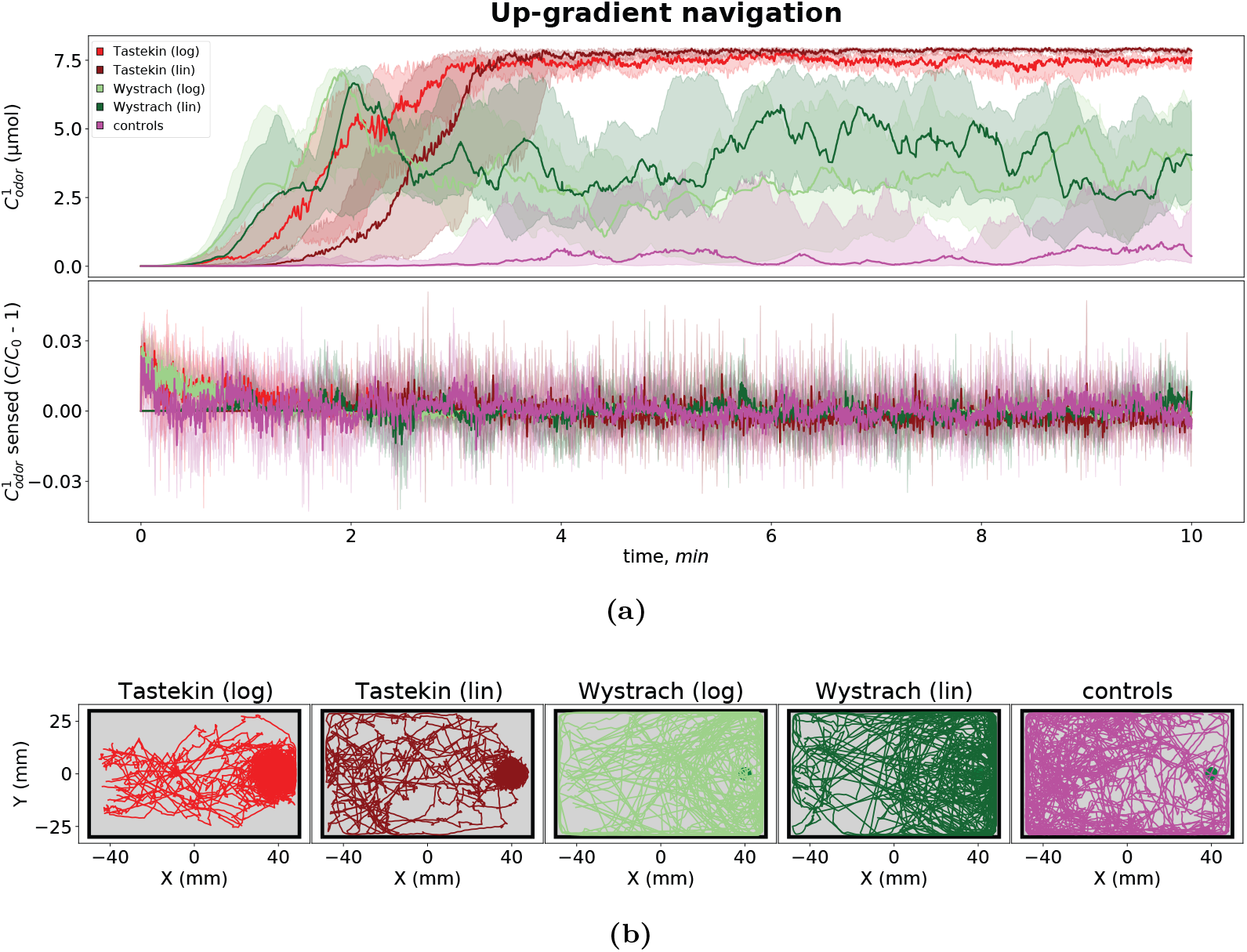
Chemotaxis towards the odor source. (a) Time plots of the absolute odor concentration (top) and perceived odor concentration change (bottom) encountered by larvae during the experiments, along with (b) the larva trajectories towards the odor source.

### Model evaluation

Evaluation of a model configuration is performed by comparing the dataset derived from the simulation of the respective virtual larva group to the empirical dataset of real larva recordings. Given that the analysis pipeline is exactly the same in both cases, the empirical and simulated datasets have exactly the same shape and format and are therefore directly comparable. As there is no established methodological framework for evaluating behavioral similarity to the real animals, we use a broad array of metrics and refrain from defining a unique global error score per model.

**Table 9.**
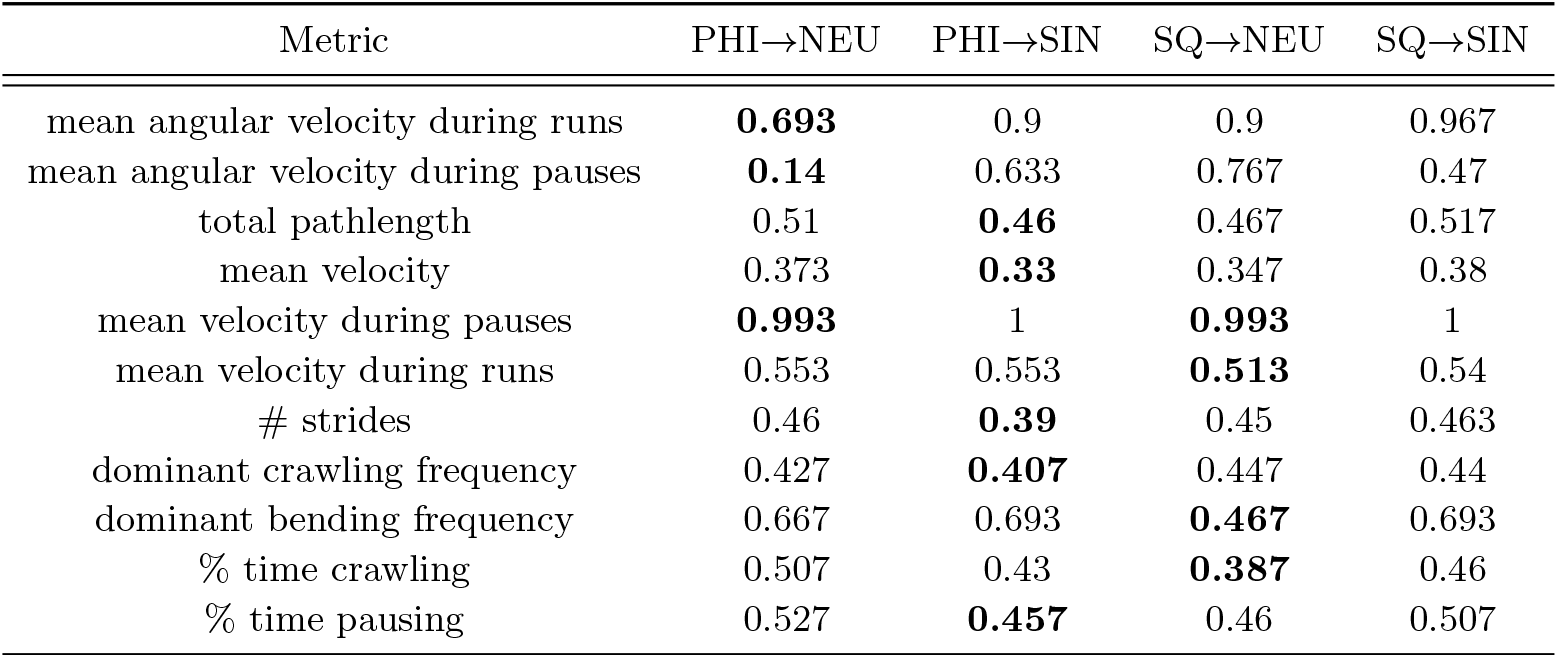
Locomotory model evaluation. Exact *KS*_*D*_ values for all endpoint-based evaluation metrics presented in Fig 12. Each row corresponds to a specific metric, and each column to one of the four evaluated model configurations. For each metric the best fitting model is highlighted in bold.

| Metric | PHI→NEU | PHI→SIN | SQ→NEU | SQ→SIN |
| --- | --- | --- | --- | --- |
| mean angular velocity during runs | **0.693** | 0.9 | 0.9 | 0.967 |
| mean angular velocity during pauses | **0.14** | 0.633 | 0.767 | 0.47 |
| total pathlength | 0.51 | **0.46** | 0.467 | 0.517 |
| mean velocity | 0.373 | **0.33** | 0.347 | 0.38 |
| mean velocity during pauses | **0.993** | 1 | **0.993** | 1 |
| mean velocity during runs | 0.553 | 0.553 | **0.513** | 0.54 |
| # strides | 0.46 | **0.39** | 0.45 | 0.463 |
| dominant crawling frequency | 0.427 | **0.407** | 0.447 | 0.44 |
| dominant bending frequency | 0.667 | 0.693 | **0.467** | 0.693 |
| % time crawling | 0.507 | 0.43 | **0.387** | 0.46 |
| % time pausing | 0.527 | **0.457** | 0.46 | 0.507 |

The evaluation metrics used are of two kinds. First, endpoint metrics where a single measurement is performed per larva, useful to summarize observations as a sum, average, variance, extrema, dominant frequency etc. Second, the timeseries of a continuous metric derived by the step-by-step simulations of individual animats, equal in shape to the number of simulation timesteps (3’ duration at 16 Hz framerate yields 2880 timesteps). In both cases the values are pooled together across the group and the error is computed as the Kolmogorov-Smyrnov distance *D*_*KS*_ between the pooled real and simulated distributions.

A summary of the evaluation process is illustrated in Fig 12. Four different model configurations have been calibrated through the pipeline described above. Specifically the models are combinations of the two turner module implementations (“NEU” for the neural oscillator, “SIN” for the sinusoidal oscillator) and the two oscillator-coupling modes (“PHI” for a continuous gaussian *c*_*CT*_ (*ϕ*_*C*_), “SQ” for a step-change of the angular suppression during a stride-cycle phase interval) The evaluation metrics are grouped into angular, spatial and temporal domains with distinct color coding for visualization purposes. The error scores of the various parameters for each of the 4 cases are min-max normalized before visualization. Exact values used for plotting are provided in Tab 9 and Tab 10.

### Individuality and variability

To test the effect of inter-individual variability in simulations, we first measured a number of endpoint parameters across a population of 200 larvae and fitted a multivariate Gaussian distribution. We select the five crawler-related parameters described in the crawler calibration section. A generated set of these parameters is adequate to completely define the crawler module. We run an evaluation simulation to compare the above-described average model to the group-level variability method. The former is therefore represented by a group of 50 identical animats while the latter by a group of 50 non-identical animats each with a parameter-set sampled from the fitted multivariate Gaussian. The results are shown in Fig 13 and Tab 11-12.

**Table 10.** Locomotory model evaluation. Exact *KS*_*D*_ values for all timeseries-derived metrics presented in Fig 12. Each row corresponds to a specific metric, and each column to one of the four evaluated model configurations. For each metric the best fitting model is highlighted in bold.

| Metric | PHI→NEU | PHI→SIN | SQ→NEU | SQ→SIN |
| --- | --- | --- | --- | --- |
| bending angle | **0.079** | 0.109 | 0.112 | 0.14 |
| angular velocity | **0.031** | 0.067 | 0.082 | 0.102 |
| angular acceleration | 0.181 | **0.166** | 0.217 | 0.181 |
| rear angular velocity | **0.089** | 0.142 | 0.145 | 0.167 |
| rear angular acceleration | **0.143** | 0.201 | 0.201 | 0.209 |
| turn-angle amplitude | 0.267 | 0.291 | **0.151** | 0.288 |
| run distance | 0.358 | **0.318** | 0.354 | 0.33 |
| # strides per run (stridechain) | 0.317 | 0.3 | 0.328 | **0.287** |
| velocity | 0.159 | 0.158 | 0.159 | **0.157** |
| acceleration | 0.086 | **0.085** | 0.086 | **0.085** |
| tortuosity over 5" | **0.241** | 0.649 | 0.303 | 0.678 |
| tortuosity over 20" | **0.224** | 0.336 | 0.351 | 0.389 |
| run duration | 0.346 | **0.318** | 0.358 | **0.318** |
| pause duration | 0.1 | 0.096 | **0.087** | 0.105 |

**Table 11.** Average vs group-variability model. Exact *KS*_*D*_ values for all endpoint-based evaluation metrics presented in Fig 13. Each row corresponds to a specific metric, and each column to one of the four evaluated model configurations. For each metric the best fitting model is highlighted in bold.

| Metric | NEU mean | NEU var | SIN mean | SIN var |
| --- | --- | --- | --- | --- |
| mean angular velocity during runs | **0.63** | 0.64 | 0.907 | 0.92 |
| mean angular velocity during pauses | **0.147** | 0.243 | 0.393 | 0.193 |
| total pathlength | 0.443 | 0.187 | 0.433 | **0.163** |
| mean velocity | 0.297 | **0.097** | 0.283 | 0.187 |
| mean velocity during pauses | 0.987 | **0.717** | 0.983 | 0.8 |
| mean velocity during runs | 0.51 | **0.337** | 0.51 | 0.36 |
| maximum dispersal in 40" | 0.713 | **0.653** | 0.713 | 0.697 |
| maximum dispersal in 60" | 0.713 | **0.653** | 0.713 | 0.697 |
| # strides | 0.303 | **0.093** | 0.283 | 0.117 |
| dominant crawling frequency | 0.427 | **0.057** | 0.437 | 0.067 |
| dominant bending frequency | **0.667** | **0.667** | 0.693 | 0.693 |
| % time crawling | 0.27 | **0.217** | 0.283 | 0.257 |
| % time pausing | 0.383 | **0.18** | 0.357 | 0.183 |

**Fig 12.**
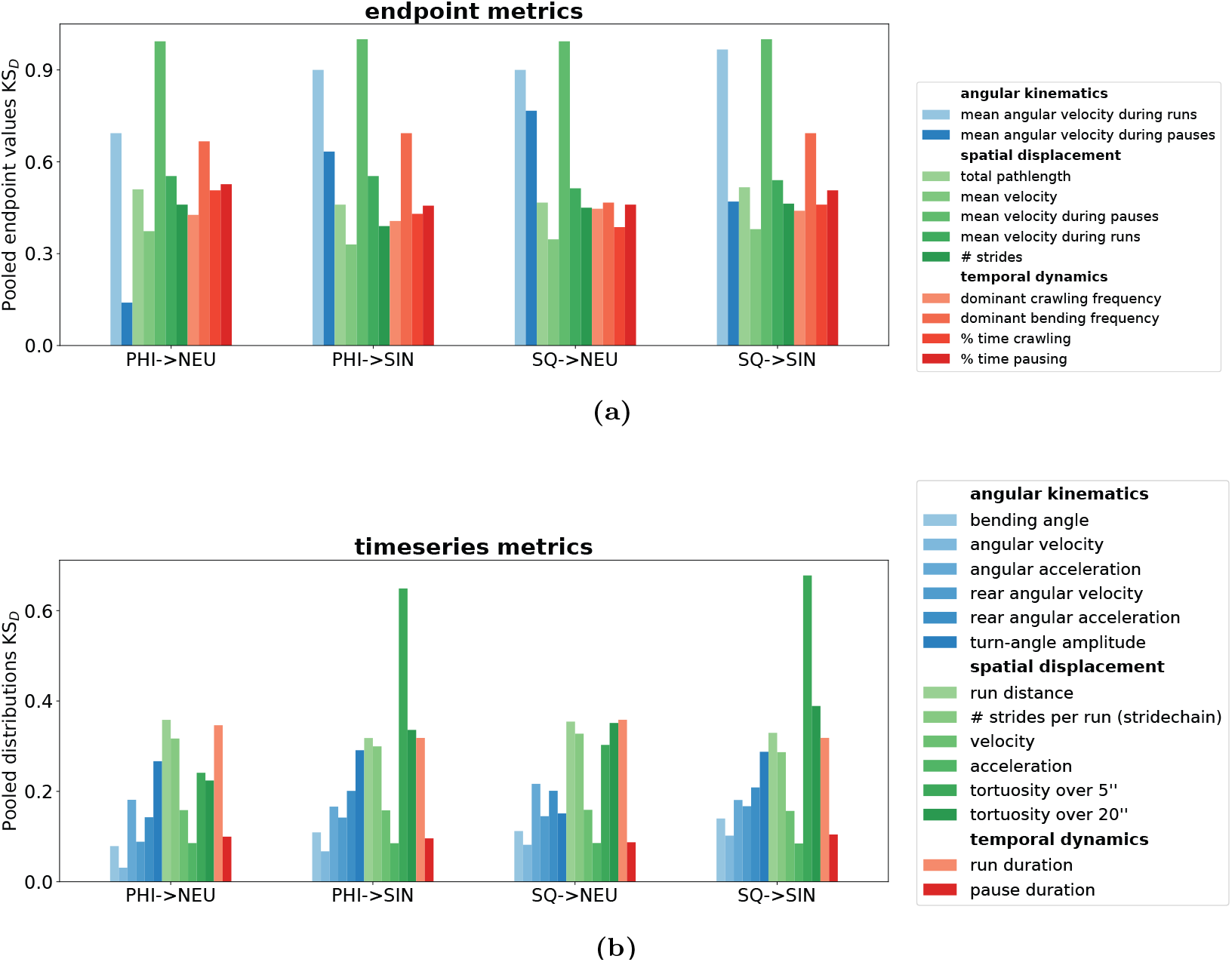
Locomotory model evaluation. Four locomotory models are evaluated against an empirical dataset of 150 larvae freely exploring a stimulus-free Petri-dish for 3 minutes. Two turner-oscillator implementations (“NEU” : neural, “SIN” : sinusoidal) are combined with two crawler-turner coupling modes (“PHI”: smooth gaussian suppression relief, “SQ” : acute step-wise suppression relief). For each model configuration 150 identical animats are placed at the exact same initial positions and with the same initial orientations as the real larvae and are simulated with the same time step as the tracked dataset over the duration of the experiment. Model evaluation is carried out over two sets of metrics: (a) single-valued endpoint measurements such as sums, averages, counts or dominant frequencies and (b) step-by-step timeseries spanning the entire experiment duration. In both cases data is pooled across the entire group and the resulting distribution’s Kolmogorov-Smyrnov distance *KS*_*D*_ to the empirical one is computed. For visualization purposes the metrics are grouped into angular, spatial and temporal domains with distinct color coding.

**Fig 13.**
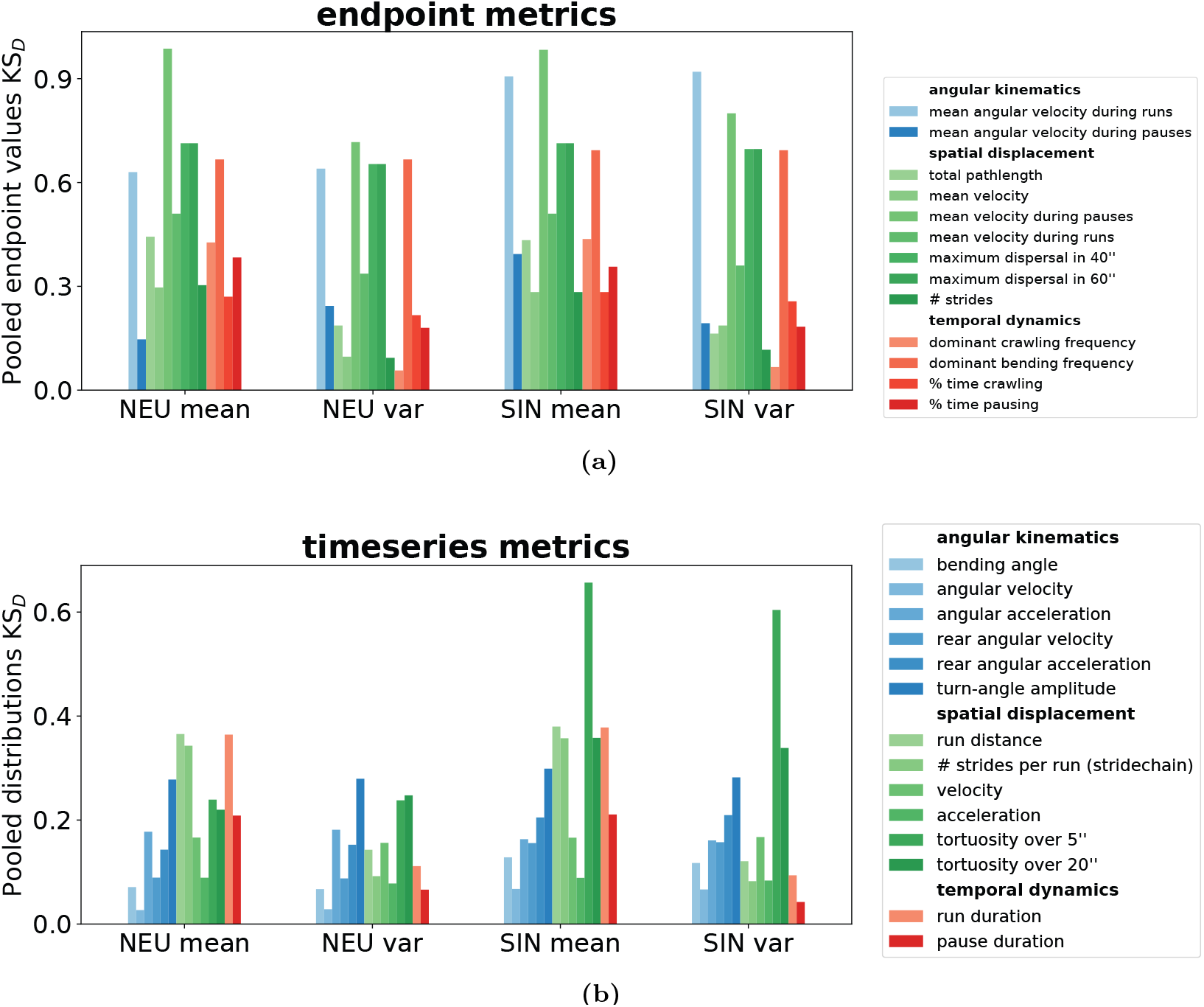
Average vs group-variability model. Two methods of generating virtual populations are compared and contrasted over each of two locomotory models already calibrated to optimally fit the average individual of the empirical dataset. A group of 50 identical animats is generated for each average model and another group of 50 non-identical animats is generated by sampling the five crawler-related parameters from a multivariate Gaussian distribution. All groups are evaluated against an empirical dataset of 150 larvae freely exploring a stimulus-free Petri-dish for 3 minutes. Model evaluation is carried out over two sets of metrics: (a) single-valued endpoint measurements such as sums, averages, counts or dominant frequencies and (b) step-by-step timeseries spanning the entire experiment duration. In both cases, data is pooled across the entire group and the resulting distribution’s Kolmogorov-Smyrnov distance *KS*_*D*_ to the empirical one is computed. For visualization purposes, the metrics are grouped into angular, spatial and temporal domains with distinct color coding.

**Table 12.** Average vs group-variability model. Exact *KS*_*D*_ values for all timeseries-derived metrics presented in Fig 13. Each row corresponds to a specific metric, and each column to one of the four evaluated model configurations. For each metric the best fitting model is highlighted in bold.

| Metric | NEU mean | NEU var | SIN mean | SIN var |
| --- | --- | --- | --- | --- |
| bending angle | 0.071 | **0.067** | 0.128 | 0.117 |
| angular velocity | **0.027** | 0.029 | 0.067 | 0.066 |
| angular acceleration | 0.178 | 0.181 | 0.163 | **0.161** |
| rear angular velocity | 0.089 | **0.088** | 0.156 | 0.157 |
| rear angular acceleration | **0.143** | 0.153 | 0.205 | 0.21 |
| turn-angle amplitude | **0.278** | 0.279 | 0.299 | 0.282 |
| run distance | 0.365 | 0.143 | 0.38 | **0.121** |
| # strides per run (stridechain) | 0.343 | 0.092 | 0.357 | **0.082** |
| velocity | 0.166 | **0.156** | 0.166 | 0.167 |
| acceleration | 0.089 | **0.078** | 0.089 | 0.084 |
| tortuosity over 5" | 0.239 | **0.238** | 0.657 | 0.604 |
| tortuosity over 20" | **0.22** | 0.248 | 0.358 | 0.339 |
| run duration | 0.364 | 0.111 | 0.378 | **0.094** |
| pause duration | 0.209 | 0.066 | 0.211 | **0.043** |

## Code availability

*Larvaworld* is available as a python package, freely distributed under the GNU General Public License 3.0. The latest version can be found at https://pypi.org/project/larvaworld/ and can be installed easily using the pip Python Installer. Code development, new features,reported issues and contributions to the project are hosted at the respective github repository (https://github.com/nawrotlab/larvaworld). Extensive documentation can be found at https://larvaworld.readthedocs.io

## Author contributions

- Conceptualization: Panagiotis Sakagiannis, Martin Paul Nawrot
- Data Curation: Tihana Jovanic
- Funding acquisition: Martin Paul Nawrot
- Methodology: Panagiotis Sakagiannis
- Software: Panagiotis Sakagiannis, Hannes Rapp
- Supervision: Martin Paul Nawrot
- Visualization: Panagiotis Sakagiannis
- Writing - Original draft: Panagiotis Sakagiannis
- Writing - Review, Editing: Panagiotis Sakagiannis, Hannes Rapp, Tijana Jovanic, Martin Paul Nawrot

## Acknowledgments

This project received funding from the Ministry of Culture and Science of the State of Northrhine Westphalia through the science network ‘iBehave’ (https://ibehave.nrw/) within the program “Netzwerke 2021”. Within the Collaborative Research in Computational Neuroscience (CRCNS) program (project ‘DrosoExpect’) funding has been received from the German Federal Ministry of Education and Research (BMBF, grant no. 01GQ2103A to M.N.), the French National Research Agency (ANR, grant no. ANR-21-NEUC-0002 to T.J.), and the United States Department of Energy (DOE, grant no. SC0021922 to B.H.S.). Additional funding has been received by T.J. from ANR-NEUROMOD (grant no. ANR-22-CE37-0027), Fédération pour la recherche sur le cerveau (FRC), and Fondation pour la Recherche Médicale: Équipe FRM EQU202303016317. We would like to thank Nick del Grosso of the iBOTS: the iBehave Open Technology Support Platform for his technical support.

## References

1. Datta SR, Anderson DJ, Branson K, Perona P, Leifer A. Computational Neuroethology : A Call to Action. Neuron. 2019;104(1):11–24. Publisher: Elsevier Inc. Available from: 10.1016/j.neuron.2019.09.038.

2. Arreguit J, Ramalingasetty ST, Ijspeert A. FARMS: Framework for Animal and Robot Modeling and Simulation. bioRxiv. 2023 Sep. Available from: http://biorxiv.org/lookup/doi/10.1101/2023.09.25.559130.

3. Zador A, Escola S, Richards B, Ölveczky B, Bengio Y, Boahen K, et al. Catalyzing next-generation Artificial Intelligence through NeuroAI. Nature Communications. 2023 Mar;14(1):1597. Available from: https://www.nature.com/articles/s41467-023-37180-x.

4. Sakagiannis P, Jürgensen AM, Nawrot MP. A behavioral architecture for realistic simulations of Drosophila larva locomotion and foraging. eLife. 2025 Apr. Available from: 10.7554/eLife.104262.1.sa3.

5. Wilson SP, Prescott TJ. Scaffolding layered control architectures through constraint closure: insights into brain evolution and development. Philosophical Transactions of the Royal Society B: Biological Sciences. 2022 Feb;377(1844). Publisher: The Royal Society. Available from: https://royalsocietypublishing.org/doi/10.1098/rstb.2020.0519.

6. Sims DW, Humphries NE, Hu N, Medan V, Berni J. Optimal searching behaviour generated intrinsically by the central pattern generator for locomotion. eLife. 2019;8:1–31. Available from: https://elifesciences.org/articles/50316.

7. Sun X, Yue S, Mangan M. A decentralised neural model explaining optimal integration of navigational strategies in insects. eLife. 2020;9:1–30.

8. Cardoso RC, Ferrando A. A Review of Agent-Based Programming for Multi-Agent Systems. Computers. 2021 Jan;10(2):16. Available from: https://www.mdpi.com/2073-431X/10/2/16.

9. An L, Grimm V, Sullivan A, Turner II BL, Malleson N, Heppenstall A, et al. Challenges, tasks, and opportunities in modeling agent-based complex systems. Ecological Modelling. 2021 Oct;457:109685. Available from: https://linkinghub.elsevier.com/retrieve/pii/S030438002100243X.

10. Foramitti J. AgentPy: A package for agent-based modeling in Python. Journal of Open Source Software. 2021 Jun;6(62):3065. Available from: https://joss.theoj.org/papers/10.21105/joss.03065.

11. Kooijman SaLM. Dynamic Energy Budget theory for metabolic organisation : Summary of concepts of the third edition. Water. 2010;365:68. ISBN: 9780521131919. Available from: http://www.pubmedcentral.nih.gov/articlerender.fcgi?artid=2981979&tool=pmcentrez&rendertype=abstract.

12. Kaun KR, Riedl CAL, Chakaborty-Chatterjee M, Belay AT, Douglas SJ, Gibbs AG, et al. Natural variation in food acquisition mediated via a Drosophila cGMP-dependent protein kinase. Journal of Experimental Biology. 2007;210(20):3547–58.

13. Thoener J, Schleyer M. Locomotion of naive Drosophila larvae. G-Node; 2021. Available from: https://doi.gin.g-node.org/10.12751/g-node.5e1ifd.

14. Gomez-Marin A, Louis M. Active sensation during orientation behavior in the Drosophila larva: More sense than luck. Current Opinion in Neurobiology. 2012;22(2):208–15. Publisher: Elsevier Ltd. Available from: 10.1016/j.conb.2011.11.008.

15. Sakagiannis PP. Larvaworld platform generated video: Wind of variable speed; 2025. Available from: https://computational-systems-neuroscience.de/wp-content/uploads/2025/03/wind_speed.mp4.

16. Sakagiannis PP. Larvaworld platform generated video: Wind-affected odorscape; 2025. Available from: https://computational-systems-neuroscience.de/wp-content/uploads/2025/03/windNodorscape.mp4.

17. Sakagiannis PP. Larvaworld platform generated video: Wind of variable direction; 2025. Available from: https://computational-systems-neuroscience.de/wp-content/uploads/2025/03/wind_direction.mp4.

18. Sakagiannis PP. Larvaworld platform generated video: Single air-puff of variable direction; 2025. Available from: https://computational-systems-neuroscience.de/wp-content/uploads/2025/03/single_air-puffs.mp4.

19. Sakagiannis PP. Larvaworld platform generated video: Repetitive air-puffs; 2025. Available from: https://computational-systems-neuroscience.de/wp-content/uploads/2025/03/repetitive_air-puffs.mp4.

20. Schumann I, Triphan T. The PEDtracker: An Automatic Staging Approach for Drosophila melanogaster Larvae. Frontiers in Behavioral Neuroscience. 2020;14.

21. Wosniack ME, Festa D, Hu N, Gjorgjieva J, Berni J. Adaptation of Drosophila larva foraging in response to changes in food resources. eLife. 2022 Dec;11:e75826. Available from: https://elifesciences.org/articles/75826.

22. Paisios E, Rjosk A, Pamir E, Schleyer M. Common microbehavioral “footprint” of two distinct classes of conditioned aversion. Learning and Memory. 2017;24(5):191–8.

23. de Tredern E, Manceau D, Blanc A, Sakagiannis P, Barre C, Sus V, et al. Feeding-state dependent neuropeptidergic modulation of reciprocally interconnected inhibitory neurons biases sensorimotor decisions in <em>Drosophila</em>. bioRxiv. 2024 Jan:2023.12.26.573306. Available from: http://biorxiv.org/content/early/2024/04/18/2023.12.26.573306.abstract.

24. Kafle T, Grub M, Sakagiannis P, Nawrot MP, Arguello JR. Evolution of temperature preference behaviour among Drosophila larvae. iScience. 2025 May:112809. Available from: https://linkinghub.elsevier.com/retrieve/pii/S2589004225010703.

25. Wystrach A, Lagogiannis K, Webb B. Continuous lateral oscillations as a core mechanism for taxis in Drosophila larvae. eLife. 2016;5.

26. Sakagiannis P, Aguilera M, Nawrot MP. A Plausible Mechanism for Drosophila Larva Intermittent Behavior. In: Vouloutsi V, Mura A, Tauber F, Speck T, Prescott TJ, Verschure PFMJ, editors. Biomimetic and Biohybrid Systems. Cham: Springer International Publishing; 2020. p. 288–99.

27. Jürgensen AM, Sakagiannis P, Schleyer M, Gerber B, Nawrot MP. Prediction error drives associative learning and conditioned behavior in a spiking model of Drosophila larva. iScience. 2024 Jan;27(1):108640. Available from: https://linkinghub.elsevier.com/retrieve/pii/S2589004223027177.

28. Sakagiannis PP. Larvaworld platform generated video: Feeding-state dependent dispersal of Drosophila larvae; 2025. Available from: https://computational-systems-neuroscience.de/wp-content/uploads/2025/03/3conditions.mp4.

29. Tastekin I, Khandelwal A, Tadres D, Fessner ND, Truman JW, Zlatic M, et al. Sensorimotor pathway controlling stopping behavior during chemotaxis in the Drosophila melanogaster larva. eLife. 2018;7:1–38.

30. Schulze A, Gomez-Marin A, Rajendran VG, Lott G, Musy M, Ahammad P, et al. Dynamical feature extraction at the sensory periphery guides chemotaxis. eLife. 2015;4(JUNE):1–52.

